# Identification and characterization of the *de novo* methyltransferases for eukaryotic N^6^-methyladenine (6mA)

**DOI:** 10.1101/2024.03.25.586193

**Authors:** Ting Cheng, Jiachen Zhang, Haicheng Li, Jinghan Diao, Wenxin Zhang, Junhua Niu, Kensuke Kataoka, Shan Gao

## Abstract

DNA N^6^-adenine methylation (6mA) is rediscovered as an important epigenetic mark in eukaryotes. We previously identified the MT-A70 family member AMT1 as the maintenance methyltransferase (MTase), but the identity of the *de novo* MTase(s) for 6mA establishment remains unidentified. The unicellular model eukaryote *Tetrahymena thermophila* offers a unique opportunity for studying *de novo* methylation: during sexual reproduction, the 6mA-free zygotic nucleus develops into 6mA-positive new macronucleus (MAC) marked by genome-wide *de novo* methylation. Here we identified that TET1 and TET2 are essential for *de novo* 6mA in the new MAC, but not for maintenance 6mA in the somatic MAC. Knockout of *TET1* and/or *TET2* not only abolished methylation on most methylatable ApT positions, but also attenuated methylation on the rest. The substantial 6mA decrease in turn altered gene expression pattern and reduced the survival rate of sexual progenies, highlighting the importance of dedicated *de novo* MTase activity.

## INTRODUCTION

The last several years have witnessed an explosively increased research interest in eukaryotic DNA N^6^-adenine methylation (6mA) ^1–7^. As a potential epigenetic mark, 6mA plays important roles in various DNA and chromatin-based molecular pathways in eukaryotes, including transcription (both activation and repression), replication, DNA damage response, and chromatin organization ^1,8–12^. It is also implicated in a plethora of physiological and pathological processes, including stress response, embryonic development, cellular physiology, plant growth and development, and tumor cell growth ^8,11,13–18^. Dissecting the molecular pathway for 6mA deposition in eukaryotes is critical for better understanding the 6mA biology.

The methylation pathway of the classical 5-methylcytosine (5mC) can be divided into two steps: establishment and maintenance. Establishment is conducted by the *de novo* methyltransferases (MTases) DNMT3A and DNMT3B ^19^, adding methyl groups to cytosines on unmethylated CpG. Maintenance is implemented by the maintenance MTase DNMT1 ^20^, which converts hemi-methylated CpG to full methylation, thus transmitting 5mC in a semi-conservative manner and preventing its dilution by DNA replication (passive demethylation) ^21^. Importantly, 5mC pattern is reset during mammalian embryonic development, with genome-wide active demethylation, followed by *de novo* methylation ^22–25^. Loss of *Dnmt3a*/*Dnmt3b* is embryonically lethal in mouse and has been linked with aberrant gene repression ^19,26,27^, underscoring the critical roles played by *de novo* DNA methylation.

6mA has been intensively studied in *Tetrahymena thermophila* (referred to as *Tetrahymena* hereafter) ^7,11,28–33^, a widely used eukaryotic model organism in laboratories ^34–36^. As featured for the binucleated ciliates, *Tetrahymena* contains a transcriptionally active macronucleus (MAC), with abundant 6mA enriched on RNA polymerase II (Pol II)-transcribed genes, and a transcriptionally silent micronucleus (MIC) with no 6mA ^11,29,30^. This pattern is consistent with the association of 6mA with transcription, as also reported in basal fungi and algae^1,37^. We and other groups have previously shown that 6mA occurs exclusively in the self-complementary ApT dinucleotides and it is maintained in a semi-conservative way by the MT-A70 family member AMT1 ^11,12,29^. All this provides definitive evidence supporting 6mA as a *bona fide* epigenetic mark.

Intriguingly, there is still 6mA in Δ*AMT1* cells, but predominantly in the form of hemi-6mApT, instead of full-6mApT as in wild-type (WT) cells ^11,29^. This strongly suggests the existence of the *de novo* MTase activity, which is largely masked by the AMT1-dependent maintenance methylation in WT cells undergoing asexual reproduction by binary fission (referred to as vegetative growth) ^38^. Most importantly, 6mA pattern also needs to be reestablished during sexual reproduction (conjugation), during which the somatic MAC is destroyed, and the germline MIC gives rise to the new MAC (Figure 1A) ^31–33^, entailing *de novo* methylation of its genomic DNA. All the evidence strongly suggests that the methylation pathway for 6mA also features two distinct stages for establishment and maintenance, in a striking parallel to 5mC.

**Figure 1.**
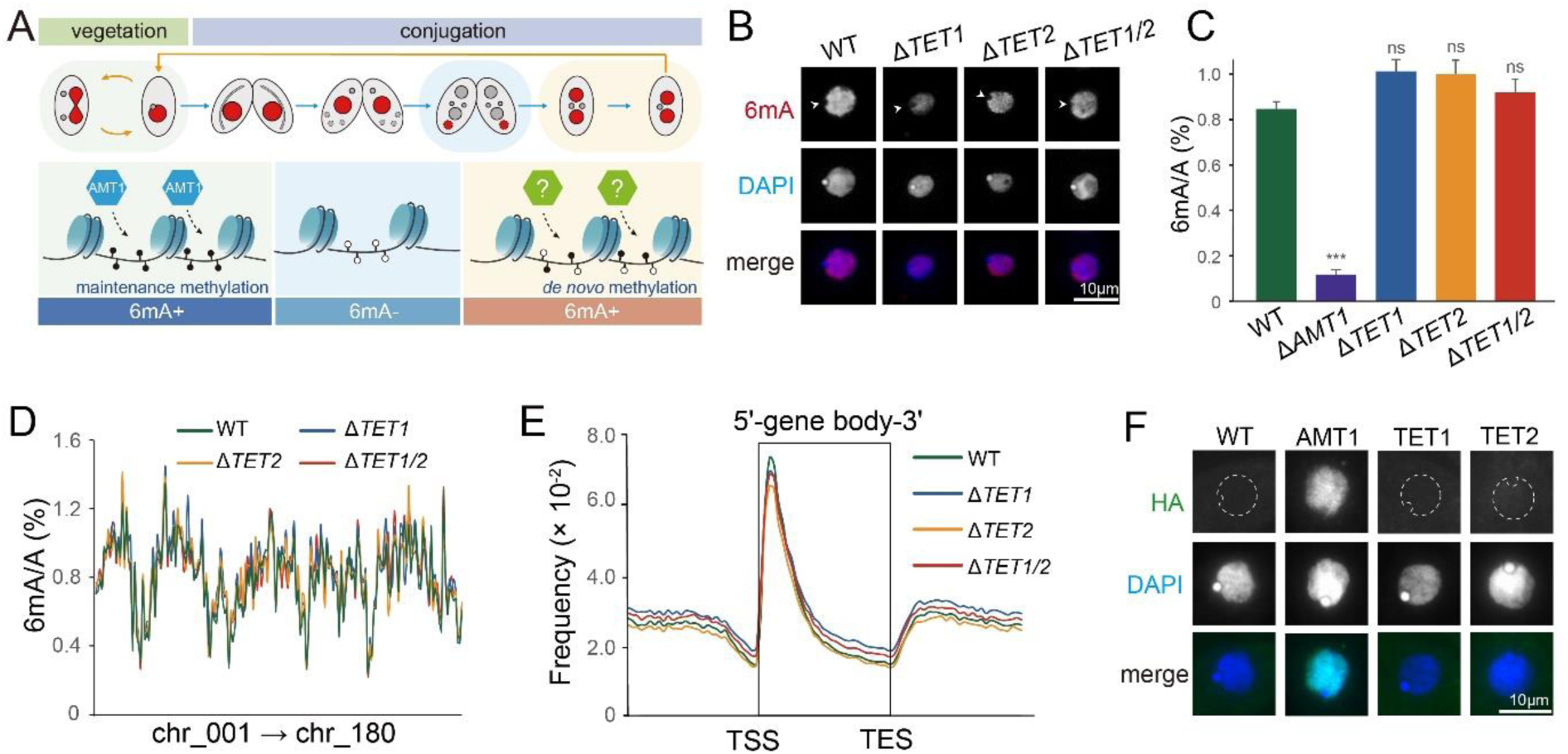
TET1 and TET2 in *Tetrahymena* are not required for maintenance 6mA. A. Top panel: schematic drawing of nuclear events and 6mA distribution during vegetation and conjugation stages (red: 6mA positive; gray: 6mA negative). Bottom panel: graphics of 6mA deposition by *de novo* and maintenance methyltransferases. B. 6mA levels were not affected in the knockout (KO) cells during the vegetative stage, as demonstrated by immunofluorescence (IF) staining. Arrowheads indicated the absence of 6mA signals in the MIC. C. Mass spectrometry (MS) analysis revealed that 6mA levels in KO cells were not significantly changed during vegetative growth. Δ*AMT1* strain was used as a control, showing a dramatic reduction in 6mA levels. D. 6mA ratios (6mA/A) of 180 non-rDNA MAC chromosomes were comparable between KO and WT cells during the vegetative stage. MAC chromosomes were arranged from left to right in accordance with their positions on the MIC chromosomes. E. 6mApT dinucleotides were enriched toward the 5’ end of gene bodies in both KO and WT cells. Genes were scaled to unit length and were extended to each side by one unit length. One unit length was divided into 30 bins and the distribution frequency of each specific position was calculated as the ratio of the 6mA amount at that position to the total 6mA amount. F. IF staining showed that TET1 and TET2 were not detectable in the somatic MAC (dotted circles) of vegetative cells. AMT1 was used as a positive control.

In this study, we utilized the *Tetrahymena* new MAC that allows us to analyze *de novo* establishment and maintenance of 6mA separately. We systematically characterized TET1 and TET2 as the candidate *de novo* 6mA MTases. We demonstrated that *de novo* 6mA in the new MAC was severely compromised in Δ*TET1*/Δ*TET2* cells. The substantial 6mA decrease in turn altered gene expression pattern and reduced the survival rate of sexual progenies, highlighting the importance of dedicated *de novo* MTase activity.

## RESULTS

### TET1 and TET2 are not required for maintenance 6mA

By performing immunofluorescence (IF) staining, we found that 6mA levels were not affected when cells lacked either one or both of TET1 and TET2 (Figures 1B, 1C and S1). This result was confirmed by mass spectrometry (MS) analysis. We then conducted Single Molecule, Real-Time Circular Consensus Sequencing (SMRT CCS), which further confirmed that neither the level nor the pattern of 6mA was altered in single or double KO cells during vegetative growth (Figures 1D and 1E, Table S2). This was most likely due to the fact that 6mA in wild-type (WT) *Tetrahymena* is almost exclusively fully methylated at the ApT dinucleotides (6mApT/6mA>99%; full-6mApT/6mApT>88%) ^29^ and efficiently maintained by the highly active AMT1 at this stage ^11,12^. In support, endogenously HA-tagged TET1 and TET2 were undetectable in the somatic MAC (Figure 1F); however, they were detected during conjugation (the sexual stage of ciliates) (Figures 2B and S1D), thus ruling out the possibility of HA tagging failure. Consistent with their dispensability, the growth rate of vegetative cells was comparable in WT and KO cells (Figure S2). Together, our results demonstrated that TET1 and TET2 are not required for maintenance 6mA in the somatic MAC.

**Figure 2.**
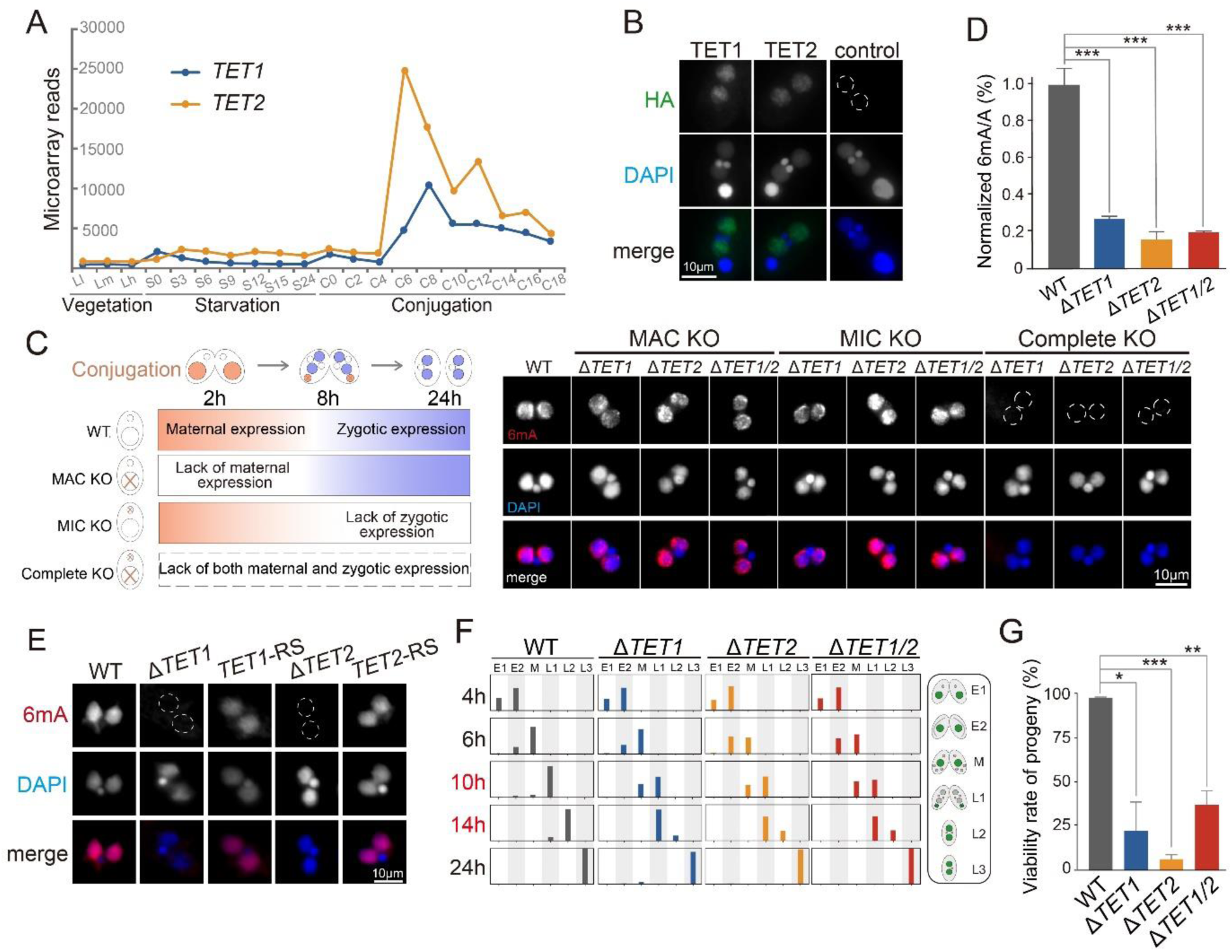
TET1 and TET2 are essential for *de novo* 6mA. A. *TET1* and *TET2* are highly expressed during new MAC formation, as demonstrated by the microarray analysis ^39^. Vegetation: three time points correspond respectively to ∼1 × 10^5^ cells/ml, ∼3.5 × 10^5^ cells/ml and 1 × 10^6^ cells/ml. Starvation: cells of ∼2 × 10^5^ cells/ml were collected at 0, 3, 6, 9, 12, 15 and 24 hours after starvation. Conjugation: equal volumes of B2086 and CU428 cells were mixed, and samples were collected at 0, 2, 4, 6, 8, 10, 12, 14, 16 and 18 hours after mixing. B. TET1 and TET2 were localized in the newly formed MAC, as demonstrated by IF staining with an anti-HA antibody. *TET1* was HA-tagged at the C-terminus and *TET2* at the N-terminus. Note the absence of HA signal in the anlagen (dotted circle) of untagged WT control cells. Cells were fixed at 12h post-mixing. C. Both somatically and zygotically expressed *TET1* and *TET2* contributed to *de novo* 6mA deposition. Left panel: expression pattern of maternal and zygotic *TET1* and *TET2* in WT and KO cells. Right panel: 6mA signals in the new MAC of MAC KO (no paternal expression), MIC KO (no zygotic expression), and complete KO cells. Note the absence of 6mA signals in the new MAC (dotted circles) of complete KO cells. Cells were fixed at 24h post-mixing. D. 6mA levels were significantly reduced in the new MAC of KO cells, as shown by MS analysis of flow-cytometry purified new MAC at 24h post-mixing. Three biological replicates were used for each strain. Data were presented as histogram (mean ± standard deviations). Student’s t-test was performed. *** P < 0.001. E. 6mA occurrence in the new MAC was restored with WT copies of *TET1* and *TET2*. The outline of the nuclei without 6mA signal were delineated with dotted circles. F. Conjugation progress in KO cells was not affected at the early stage but was delayed during new MAC development, especially at 10∼14h post-mixing. Mating cells of Δ*TET1* (IV × V), Δ*TET2* (II × IV), Δ*TET1*/*2* (II × IV), and WT (SB210 × CU428) at each time point were classified into different developmental stages according to their nuclear morphology: pre-meiosis (E1), meiosis (E2), mitosis (M), new MAC development (L1), pair separation (L2), and one of the new MIC degraded (L3). n>200 for each strain. n represented the number of single cells. G. The viability rate of KO progenies was significantly reduced compared to that of WT progenies. Data were presented as histogram (mean ± standard deviations). Student’s t-test was performed. * P < 0.05, ** P < 0.01, *** P < 0.001

### TET1 and TET2 are required for *de novo* 6mA

To determine whether TET1 and TET2 are responsible for *de novo* 6mA methylation, we first examined their expression patterns. Both *TET1* and *TET2* showed a two-peak profile, one at early conjugation, and the other at late conjugation corresponding to the stage of new MAC development (Figure 2A)^39^. Consistent with their expression profiles, we detected the endogenously tagged TET1 and TET2 first localizing in the parental MAC and then in the newly formed MAC (Figures 2B and S1D).

During the stage of new MAC development, 6mA in WT cells exclusively occurred in the new MAC of exconjugants (when two progenies separate) (Figure 2C). Interestingly, in MAC KO cells only lacking the maternal transcripts of *TET1* and *TET2*, 6mA signals remained unchanged (Figure 2C). Similarly, 6mA signal intensity was not affected in MIC KO cells with the exclusive absence of the zygotic transcripts (Figure 2C). Only in complete KO cells where both maternal and zygotic transcripts were eliminated, did 6mA signals become undetectable (Figure 2C). This dramatic reduction of 6mA was corroborated by the MS analysis conducted on flow cytometry-purified new MACs of complete KO cells (Figure 2D). We therefore conclude that the deposition of *de novo* 6mA is attributable to both maternally and zygotically expressed *TET1* and *TET2*.

To test whether the 6mA absence in the new MAC was solely attributable to the lack of *TET1* or *TET2*, we introduced WT *TET1* or *TET2* to their endogenous MAC locus of corresponding KO cells (Figures S4A-B and S4D-E). Western blot analysis confirmed that TET1 and TET2 were restored to WT levels in rescued cells (Figures S4C and S4E). More importantly, 6mA signal intensity was restored to WT level (Figure 2E). We also mutated the potential catalytic residues in TET1 and confirmed that the intrinsic enzymatic activity of TET1 underpinned its function for 6mA methylation.

Next, we tested the effect of 6mA deficiency on conjugation. Compared to WT cells, the early conjugation progress was not affected in cells lacking *TET1* and/or *TET2* (Figure 2F). However, late conjugation progress, especially during 10-14 hours post-mixing, suffered major delays in both single and double KO cells (Figure 2F). Although such delays could be subsequently mitigated, possibly due to the compensational effect of AMT1 (Figure 6), viability rates of KO progenies were significantly lower than WT counterparts (Figures 2G and S5, Table S3), underscoring the functional significance of TET1/TET2 and *de novo* 6mA for conjugation progression and progeny development.

It should be noted that we observed no synergistic or additive effect for the double KO of *TET1* and *TET2*, compared to single KO. Moreover, the global transcription profile of the conjugation progenies for single and double KO cells were more similar to each other than to WT control cells (Figure S6), strongly indicating that TET1 and TET2 function cooperatively rather than independently.

Together, these results strongly argued that TET1 and TET2 are essential for the *de novo* establishment of 6mA in the new MAC, as well as for the cell fitness.

### 6mA landscape in the new MAC

We next performed SMRT CCS of flow-cytometry purified new MACs from both WT and KO cells. After applying a stringent threshold and filtering out abnormal DNA molecules (see more details in Methods and Materials) ^29^, 1,627,095, 1,874,423, 1,974,209, and 1,153,747 high-confidence single molecules for WT, Δ*TET1,* Δ*TET2* and Δ*TET1/2* cells were used for further analysis (Table S4).

WT new MAC (WT-C24) exhibited striking similarity to WT somatic MAC during the vegetative growth (WT-veg) ^29^. ***1)*** 6mA occurred almost exclusively at the ApT dinucleotides in the new MAC (6mApT/6mA>99%), consistent with WT-veg (>99%) (Figures 3A and S7A). 6mA in non-ApT dinucleotides (ApC/ApG/ApA) were called at low levels close to the background noise using the same threshold (Figures 3A and S7B). ***2)*** 6mApT represented 2.25% of all mapped ApT sites (6mApT/ApT) (vs. 2.03% in WT-veg) (Table 1). Correspondingly, 6mA level (6mA/A) was comparable between two samples (0.82% in WT-C24 vs. 0.74% in WT-veg) (Table 1). ***3)*** There were significant overlaps in ApT positions between WT-C24 and WT-veg; most ApT positions methylated in WT-veg were also methylated in WT-C24 (Figure 3B). When 6mApT positions were divided into ten quantiles according to their penetrations (ratio between the number of 6mA sites and all adenine sites in all SMRT CCS reads) in WT-veg, 6mApT in WT-C24 displayed a similar distribution in each quantile; sites of high penetration in WT-veg had a higher tendency to be also highly-methylated in WT-C24 (Figure 3C). ***4***) 6mA in both samples were enriched at gene bodies, towards the 5’ end and downstream of the transcription start sites (TSS) (Figure 3D) ^11,28^. ***5)*** 6mA levels at individual genes showed strong correlation between WT-C24 and WT-veg (Figure 3E, Spearman’s correlation coefficient=0.93, k=0.99). Collectively, these connections matched with the scenario that 6mA landscape was nearly established in the new MAC, before cells reentered the vegetative growth stage^31^.

**Figure 3.**
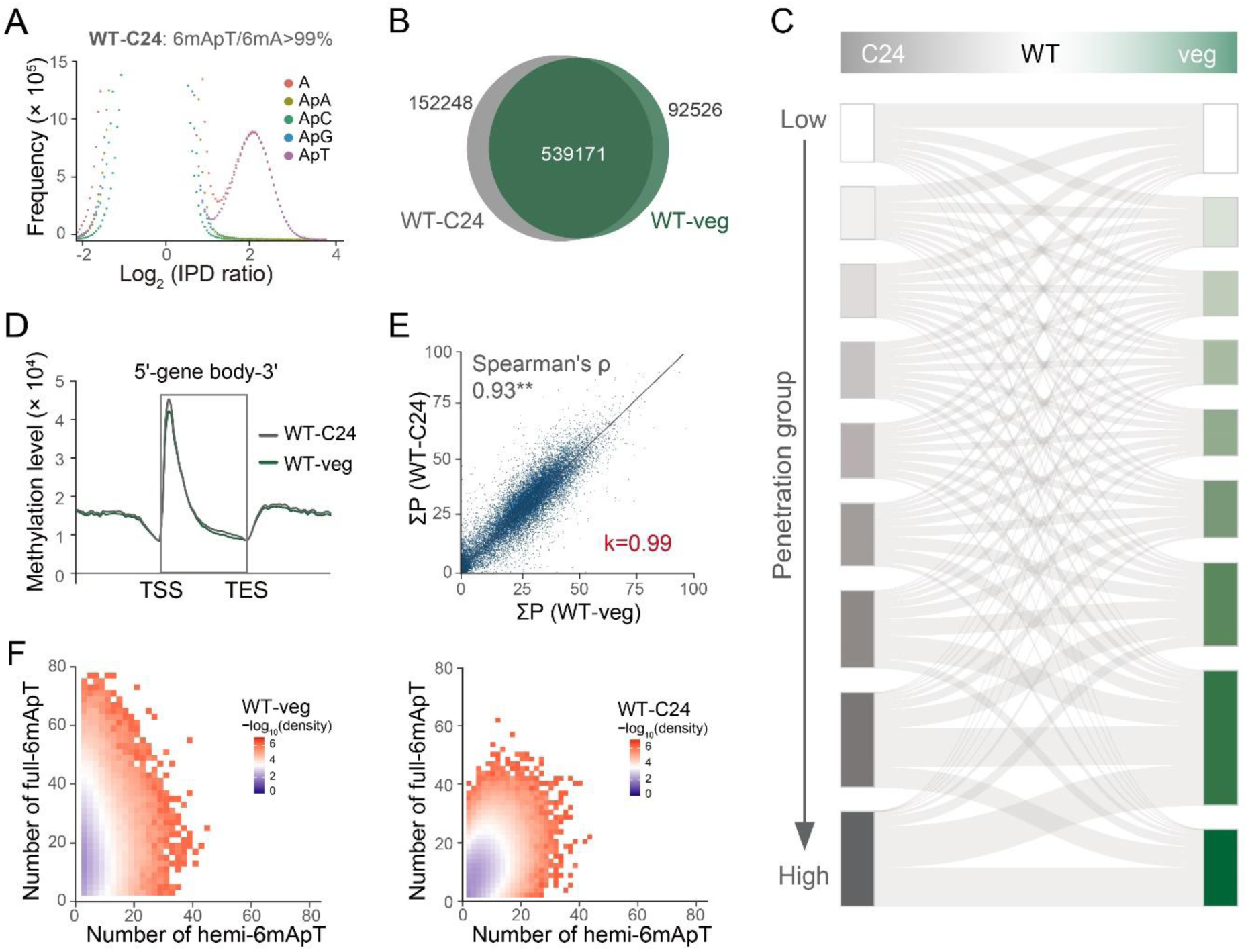
The characteristics of 6mA in WT-C24 were similar to those in WT-veg. A. 6mA in the new MAC of WT conjugative cells (WT-C24) exhibited a preference for ApT dinucleotides. The IPD ratios distribution of total adenine and all kinds of adenine-dinucleotides were plotted. The IPD ratios of adenine showed a significant bimodal distribution and the right peak with higher IPD ratios represented 6mA. Adenines and ApT dinucleotides displayed similar IPD ratio distribution features, while non-ApT dinucleotides only had the left IPD ratios peak, corresponding to unmethylated adenines. B. Venn diagram showed a strong overlap of 6mApT positions between the somatic MAC of WT vegetative cells (WT-veg) and WT-C24 (representation factor > 1, P-value < 0.05). 6mApT positions with sufficient 6mApT coverage (≥ 10×) were selected for analysis. C. Sankey diagram showed penetration profiles of 6mApT positions in WT-C24 and WT-veg. 6mApT positions were divided into 10 groups according to their penetration, which was defined as “the ratio between the number of 6mA sites and all adenine sites at a specific position”. Only 6mApT positions with a penetration > 0.1 were included in the analysis. D. 6mApT was enriched toward the 5’ end of gene bodies. Genes were scaled to a unit length and were extended by one unit length on each side. One unit length was divided into 30 bins and the sum of penetration within each bin represented the degree of 6mApT enrichment. E. 6mA levels of individual genes were nearly identical in WT-C24 and WT-veg. 6mA level of a specific gene was calculated as “the sum of penetration of all 6mApT positions (∑P)”. F. The proportion of full-6mApT sites were lower in WT-C24 than in WT-C24, demonstrated at the single molecule level. Hemi-6mApT (Y-axis) and full-6mApT (X-axis) dinucleotides were counted for each single molecule. The number of single molecules per bin was divided by the total number of single molecules in each sample for normalization.

**Table 1.** SMRT CCS results for flow-cytometry purified new MAC at 24h post-mixing.

| Single molecule | WT-C24 | | $\Delta TET1$-C24 | | $\Delta TET2$-C24 | | $\Delta TET1/2$-C24 | | WT-Veg* | | $\Delta AMT1$-Veg* | |
| --- | --- | --- | --- | --- | --- | --- | --- | --- | --- | --- | --- | --- |
|  | Number | Percentage (%) | Number | Percentage (%) | Number | Percentage (%) | Number | Percentage (%) | Number | Percentage (%) | Number | Percentage (%) |
| A sites | 2,650,998,266 | - | 4,562,894,840 | - | 4,138,411,182 | - | 2,963,324,787 | - | 2,430,618,393 | - | 1,730,640,574 | - |
| 6mA sites | 21,821,071 | 0.82 | 5,524,473 | 0.12 | 4,296,775 | 0.10 | 3,977,697 | 0.13 | 17,889,360 | 0.74 | 3,443,531 | 0.20 |
| ApT sites | 917,555,698 | - | 1,573,597,700 | - | 1,429,134,888 | - | 1,021,233,408 | - | 823,903,428 | - | 645,000,722 | - |
| 6mApT sites | 20,651,604 | 2.25 | 4,750,706 | 0.30 | 3,502,851 | 0.25 | 3,275,705 | 0.32 | 16,765,614 | 2.03 | 3,420,511 | 0.53 |
| Full | 15,531,852 | 75.21 | 3,828,384 | 80.59 | 2,758,540 | 78.75 | 2,623,536 | 80.09 | 14,811,496 | 88.35 | 82,362 | 2.40 |
| Hemi-C | 2,569,180 | 12.44 | 461,348 | 9.71 | 372,416 | 10.63 | 326,384 | 9.96 | 974.677 | 5.81 | 1,670,284 | 48.83 |
| Hemi-W | 2,550,572 | 12.35 | 460,974 | 9.70 | 371,895 | 10.62 | 325,785 | 9.95 | 979,261 | 5.84 | 1,667.685 | 48.77 |
| Total 6mApT | 20,651,604 | - | 4,750,706 | - | 3,502,851 | - | 3,275,705 | - | 16.765.614 | - | 3.420.511 | - |

\* Data from Sheng et al., 2023

Despite these connections, there were distinctions between 6mA in WT-C24 and WT-veg. First, the percentage of fully-methylated 6mA was lower in WT-C24 than in WT-veg (75.21% vs. 88.35%) (Figure S7C, Table 1), a phenomenon commonly observed at the individual single-molecule level (Figure 3F). This might be partially attributable to the low activity of AMT1 during the new MAC development (Figure 6D). Second, some highly-methylated sites in WT-C24 were not transmitted and maintained in WT-veg (Figure 3C), which were likely to be independent of AMT1.

### TET1 and TET2 are required for the proper establishment of the 6mA landscape

In contrast to WT cells (WT-C24), 6mApT/ApT ratio in the new MAC was dramatically reduced to 0.30%, 0.25% and 0.32% in Δ*TET1,* Δ*TET2* and Δ*TET1/2* cells, respectively (Table 1). Correspondingly, 6mA level (6mA/A) in KO cells was remarkably lower than that in WT cells in all 180 non-rDNA chromosomes (0.12%, 0.10% and 0.13% in Δ*TET1,* Δ*TET2* and Δ*TET1/2* vs. 0.82% in WT cells) (Figure 4A, Table 1), consistent with the IF staining and MS results (Figures 2C and 2D). This reduction is even more dramatic than cells depleting *AMT1* (Δ*AMT1*, 6mA/A=0.20%) ^11,29^.

**Figure 4.**
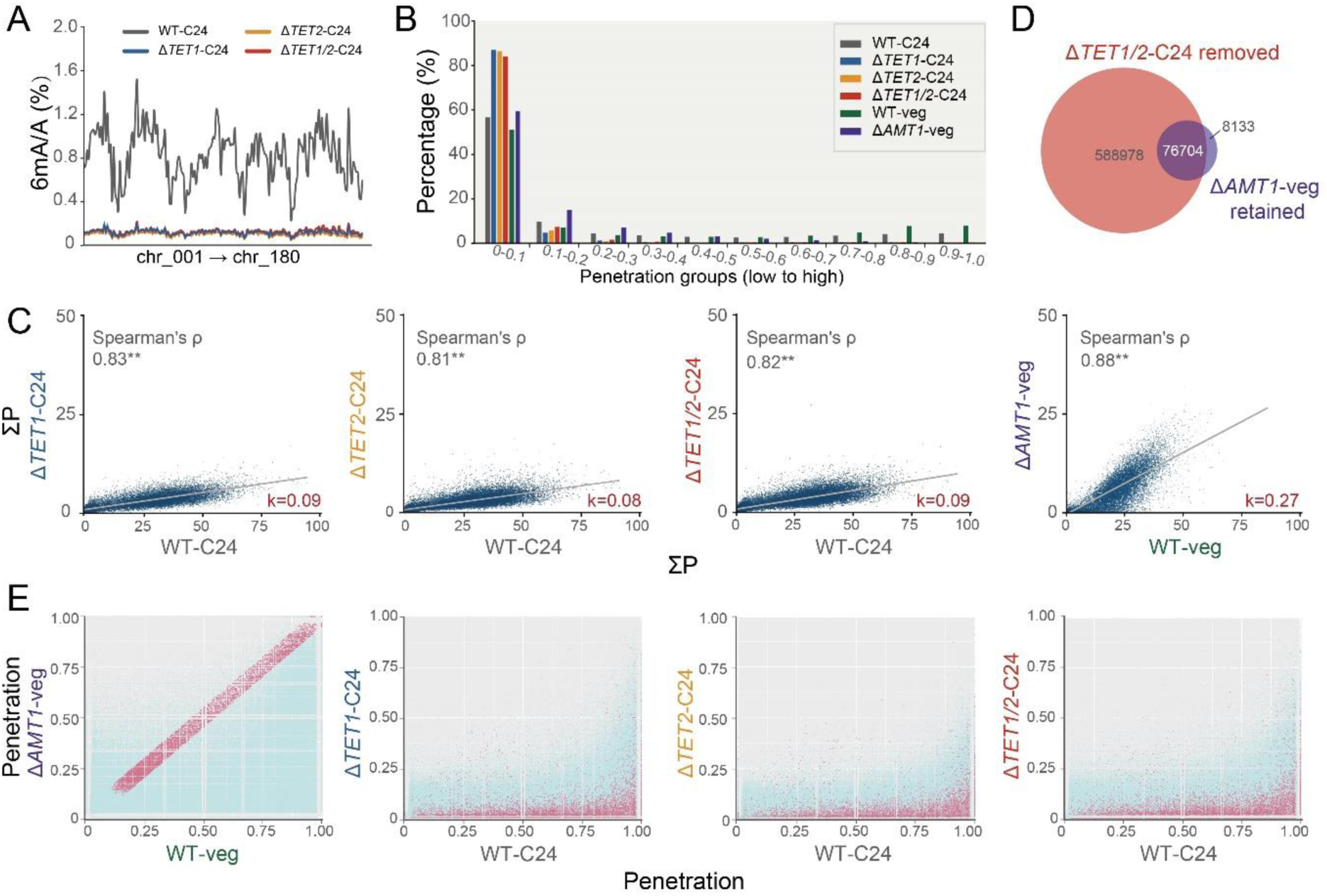
6mA establishment was severely impaired in *TET1* and/or *TET2* KO cells. A. 6mA levels, defined as the percentage of methylated A among all A (6mA/A), exhibited a global decrease across all 180 non-rDNA chromosomes in Δ*TET1*, Δ*TET2*, and Δ*TET1*/*2* cells at 24h post-mixing (C24). Chromosome numbering followed the assignment outlined in Sheng et al. (2020). B. 6mApT positions with high penetration in WT-C24 cells were depleted in Δ*TET1*/*2*-C24. 6mApT positions were divided into 10 groups according to their penetration, from low to high. The extent of reduction was even more pronounced than observed in vegetative Δ*AMT1* cells (Δ*AMT1*-veg). C. 6mA levels of individual genes were dramatically reduced in Δ*TET1*-C24, Δ*TET2*-C24, Δ*TET1/2*-C24 as well as in Δ*AMT1*-veg. 6mA level of a specific gene was calculated as “the sum of penetration of all 6mApT positions (∑P)”. D. Venn diagram illustrated that the vast majority of the retained 6mA sites in Δ*AMT1* cells were unmethylated in the new MAC of Δ*TET1/2* cells. Δ*TET1*/*2*-C24 removed sites were as defined as “positions with less than 10× coverage of 6mApT in Δ*TET1*/*2*-C24”. Δ*AMT1*-veg retained sites were defined as “positions with more than 10× coverage of 6mApT in Δ*AMT1*-veg”. E. The penetration of AMT1-independent 6mApT positions was more significantly reduced in Δ*TET1*/*2* cells. AMT1-independent 6mApT positions, depicted as red points, were defined as having a penetration difference ≤ ±0.05 between WT-veg and Δ*AMT1*-veg, with 6mApT coverage ≥ 5× in both cells. The reduction in penetration of AMT1-independent 6mApT positions were more pronounced than the rest majority of 6mApT positions in Δ*TET1*-C24, Δ*TET2*-C24 and Δ*TET1*/*2*-C24.

Many ApT positions were methylated in WT cells but not in *TET1* and/or *TET2* KO cells (Figure 5), with a small proportion of overlapped sites (3.74% of WT 6mA sites) (Figures 6A and S8A). Along with the global reduction of methylated 6mA sites (Figure 4A, Table 1), the penetration of individual 6mA sites was generally much lower in KO cells; 6mA positions of high penetration were especially depleted (Figure 4B and Figure 5). At individual genes, 6mA levels were also substantially reduced, more dramatic than in Δ*AMT1* cells (Figure 4C). Together, these results strongly argued for the essential role of TET1 and TET2 to establish the proper 6mA landscape in the new MAC.

**Figure 5.**
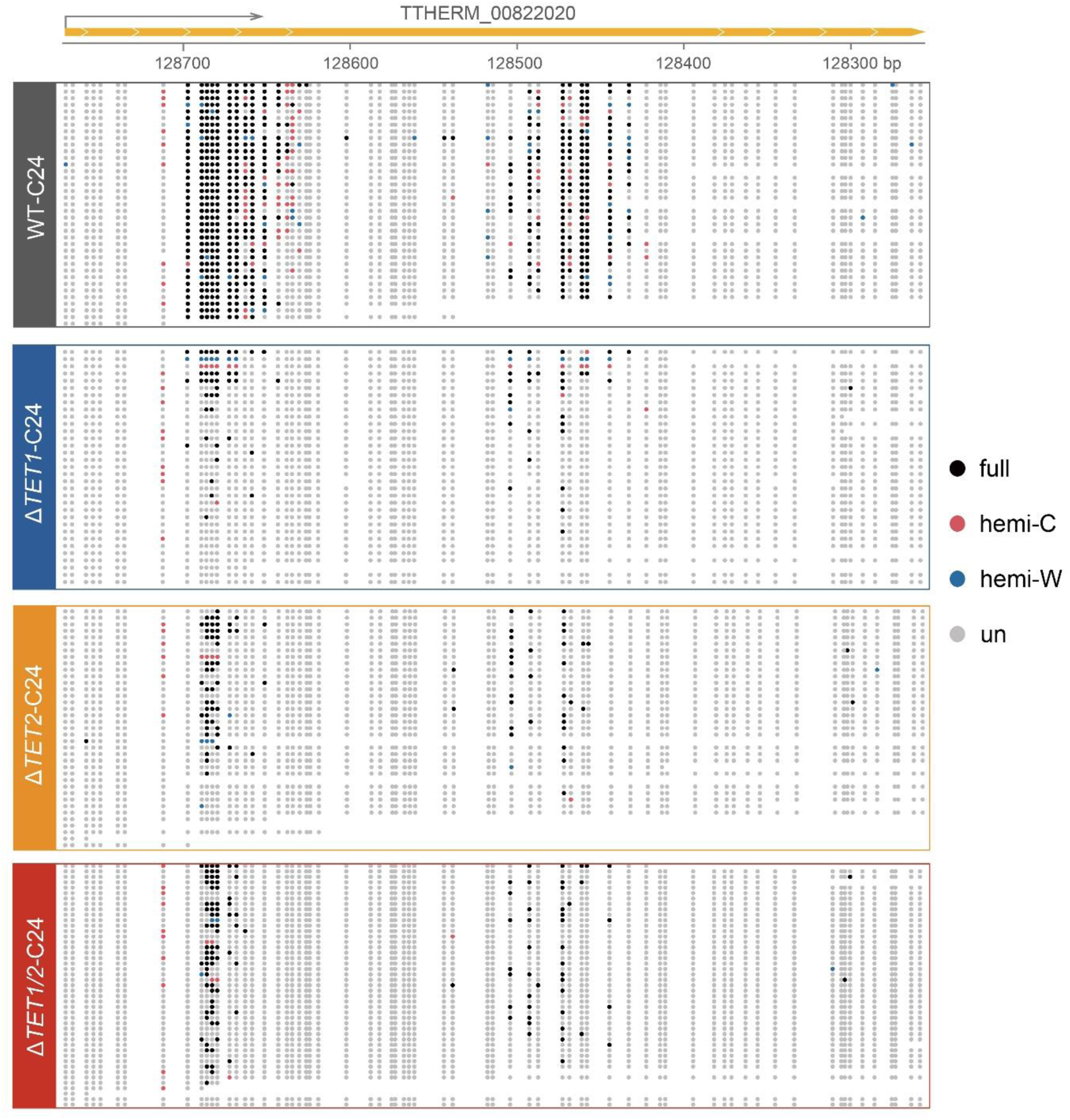
Typical DNA single molecules revealed that a substantial proportion of highly-methylated 6mApT sites in WT cells were either unmethylated or lowly-methylated in KO cells. C24 represented new MACs collected at 24h post-mixing by flow cytometry.

To more directly interrogate the features of TET1/TET2-deposited 6mA, we first selected 6mA sites remaining in Δ*AMT1* cells that were presumably methylated by TET1 and TET2. Intriguingly, 90.4% of these sites were unmethylated in the new MAC of cells lacking *TET1* and *TET2* (Figure 4D). Next, to minimize the effect of AMT1, we focused on sites displaying comparable penetrations in WT (WT-veg) and in Δ*AMT1* cells (Δ*AMT1*-veg) (Figure 4E) ^29^. These identified sites were designated as AMT1-independent, enabling a targeted evaluation of their reliance on TET1 and TET2. Penetration of these selected sites was substantially lower in KO cells compared to WT cells, consistent with the trend observed at other sites (Figure 4E). Remarkably, their penetration was much more diminished to zero in KO cells than the rest sites (Figure 4E), highlighting the significant contribution of TET1 and TET2 in their modification. Given that these sites are predominantly in hemi-methylated state in Δ*AMT1* cells ^29^, we postulated that TET1/TET2-deposited 6mA are hemi-methylated.

### AMT1-catalyzed 6mA in the new MAC without TET1/TET2

Albeit at a notably lower level (Figure 4B and Figure 5), the vast majority of remaining 6mA sites in the new MAC of KO cells were also methylated in WT cells (Figures 6A and S8A). The methylation pattern of these remaining 6mA in KO cells was reminiscent of that in WT cells; they still occurred at the ApT dinucleotides towards the 5’ end of the gene body (Figure 5 and Figures 6B-C)^11^. Moreover, the full/hemi ratio (defined as ratio between full-6mApT and hemi-6mApT) in KO cells remained similar to that in WT cells (Table 1). Full-6mApT in KO cells was likely not generated by the random combination of two independent hemi-methylation events, as the penetration strand bias of 6mA in KO cells converged to zero along with the increase of 6mA coverage (Figure S7D) ^29^.

The similar 6mA pattern in the new MAC of both WT and KO cells was presumably the product of AMT1. We favored the scenario that in the absence of TET1 and TET2, AMT1 catalyzed *de novo* methylation on a limited number of ApT sites and converted a large majority of them into full methylation ^29^. Indeed, AMT1 was localized into the new MAC ^11^ and its protein level in the new MAC was comparatively lower than in the somatic MAC of conjugation progenies after refeeding (Figure 6D), partially explaining the lower full/hemi ratio in the new MAC (Table 1). We also noticed that AMT1 protein level was upregulated in KO cells than in WT cells during late conjugation and after refeeding (Figure 6D), which presumably contributed to 6mA recovery (Figure 6E).

**Figure 6.**
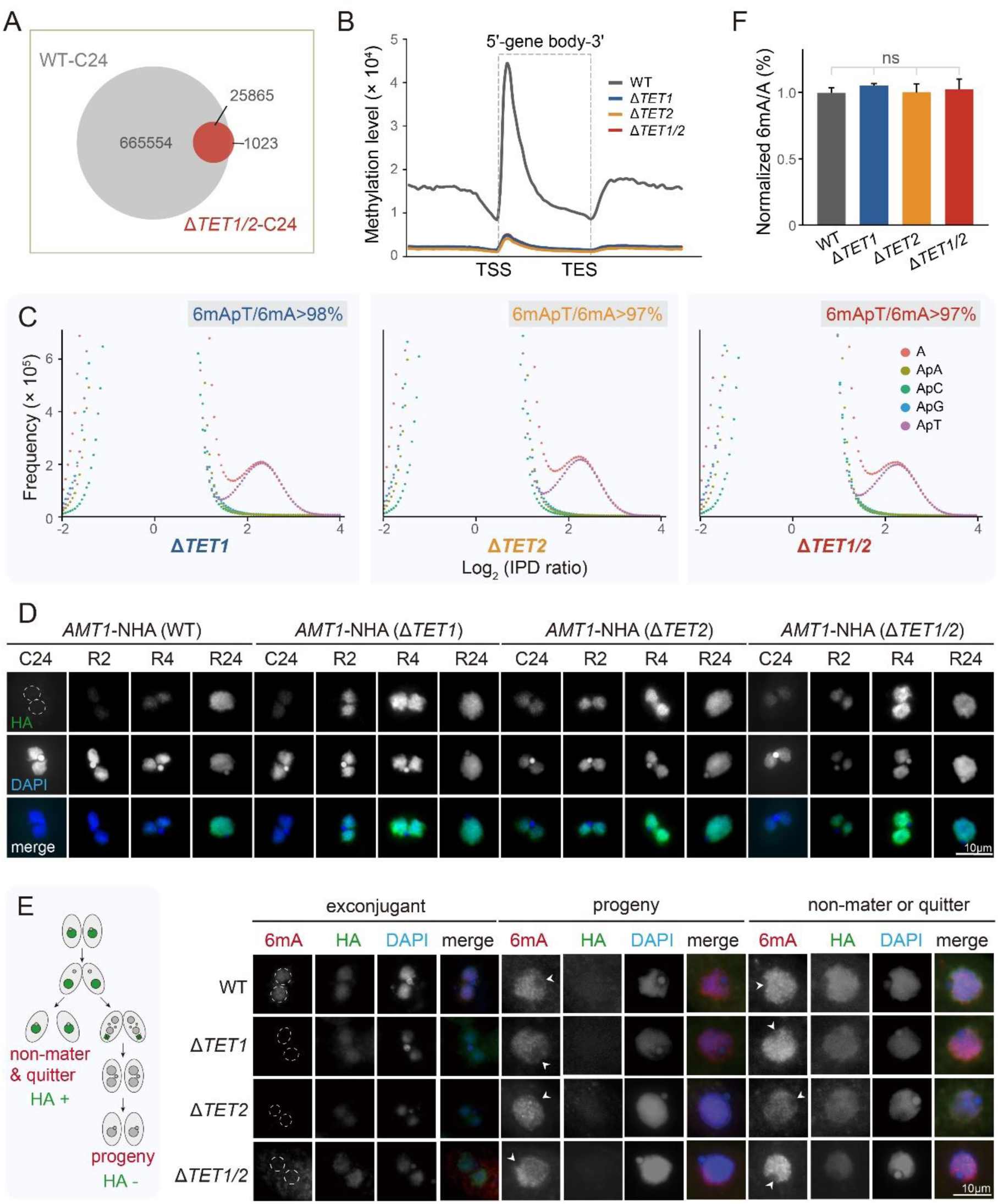
The methylation pattern of the remaining 6mA in the new MAC of KO cells was attributed to AMT1. A. Venn diagram illustrated the overlap between methylated 6mA sites in WT-C24 and Δ*TET1*/*2*-C24. The vast majority of 6mApT positions in Δ*TET1/2*cells was also methylated in WT cells. The same cutoff (6mApT coverage ≥10×) was applied to both samples. B. 6mApT in KO cells still exhibited an enrichment towards the 5’ end of gene bodies, although with a much decreased extent of enrichment. C. 6mA in KO cells displayed a preference for ApT dinucleotides. Note that the bimodal distributions of both adenines (A) and ApT dinucleotides (ApT) were less obvious in KO cells, attributable to the highly reduced number of 6mApT sites. D. AMT1 levels were upregulated in KO cells both at late conjugation and after refeeding, demonstrated by IF staining. AMT1 was N-terminally HA-tagged in both MAC and MIC endogenous loci. The outline of the nuclei without HA signal were delineated with dotted circles. C24: at 24h post-mixing. R2, R4, R24: 2h, 4h, and 24h after refeeding. E. 6mA signal intensity was restored after refeeding in exconjugants of KO cells. Left panel: schematic diagram showing that the *RPB3*-CHA construct was transformed into somatic MACs to distinguish progeny (HA negative) from non-mater or quitter (HA positive). Right panel: 6mA occurrence in KO exconjugants before refeeding was abolished, but was restored after refeeding. New MACs were circled with the white dotted line and the arrowheads indicated the absence of 6mA signals in MICs. F. MS analysis demonstrated that 6mA levels in KO progenies were comparable to their WT counterparts. Two biological replicates were performed for each strain.

To monitor the 6mA recovery, we introduced a hemagglutinin (HA) tag to the endogenous *RPB3* locus in the somatic MAC, to distinguish true progeny (HA negative, somatic MAC degraded) from non-maters and quitters (HA positive, somatic MAC retained) (Figure 6E, left panel). In the presence of AMT1, 6mA level in progenies of KO cells could be restored to that of WT cells (Figure 6E, right panel), at as early as ten hours after refeeding (Figure S8B). MS analysis also showed that 6mA level in the conjugation progenies of KO cells were comparable to their WT counterparts (Figure 6F).

## DISCUSSION

### TET1 and TET2 are *de novo* 6mA methyltransferases

We and others have demonstrated that AMT1 functioned as the maintenance MTase, displaying a higher enzymatic activity towards the hemi-methylated ApT than unmethylated ApT ^11,12,29^. Notably, AMT1 also had a comparatively lower *de novo* MTase activity, being able to catalyze methylation on unmethylated substrates, demonstrated by *in vitro* assay ^12,29^. However, when Δ*AMT1* cells mated with one another, the majority of mating pairs could not progress to the new MAC stage ^11^, thus constraining our capacity to investigate the *de novo* deposition of 6mA during new MAC formation. Nevertheless, among the rare Δ*AMT1* survivors that succeeded to enter the new MAC stage, *AMT1* deletion caused dramatic 6mA hypomethylation but not a complete loss^11^. This observation strongly suggested the involvement of alternative MTases in *de nov*o 6mA deposition ^11,12,29^.

In this study, we demonstrated that knockout of *TET1* and/or *TET2* not only abolished methylation on a large majority of ApT sites, but also attenuated the methylated ApT sites to reach high penetration. The dramatic 6mA decrease in turn had a profound impact on the survival of conjugation progenies. The features of TET1/TET2-catalyzed 6mA could be partially deduced from the remaining sites in Δ*AMT1* cells, with the preferential accumulation on ApT dinucleotides, the 5’ end of gene body, and linker DNA regions, and most importantly, in hemi-methylated forms ^11,29^.

In WT cells, the establishment of 6mA landscape during late conjugation was the coordinated work of TET1/ TET2 and AMT1 (Figure 7). During the transition from 6mA-free zygotic MIC to 6mA-positive new MAC ^31^, TET1 and TET2 catalyzed unmethylated adenines into hemi-6mA, which was further converted into full-6mA by AMT1. In *TET1* and/or *TET2* KO cells, however, the impaired un-to-full conversion became the limiting step for AMT1 function. Instead, AMT1 conducted a two-step catalyzation on its own, un-to-hemi and then hemi-to-full. The former reaction was not effective enough, with only 10-20% of WT 6mApT sites being methylated at low penetrations, highlighting the indispensability of TET1 and TET2 for the establishment of proper 6mA landscape. It also provided strong *in vivo* evidence for the ability of AMT1 to target unmethylated DNA. The latter reaction was comparatively efficient, resulting in the typical footprint of AMT1 such as the predominance of full-6mA and the enrichment at the 5’ end of gene bodies ^11,28,29^. It also suggested that AMT1, or even TET1 and TET2, are interacting with other epigenetic factors during new MAC development, to achieve their genomic targeting. Those factors could still be H3K4me3 and H2A.Z, as demonstrated in vegetative cells^11,28^, which are already abundant in the new MAC at this stage ^40,41^.

**Figure 7.**
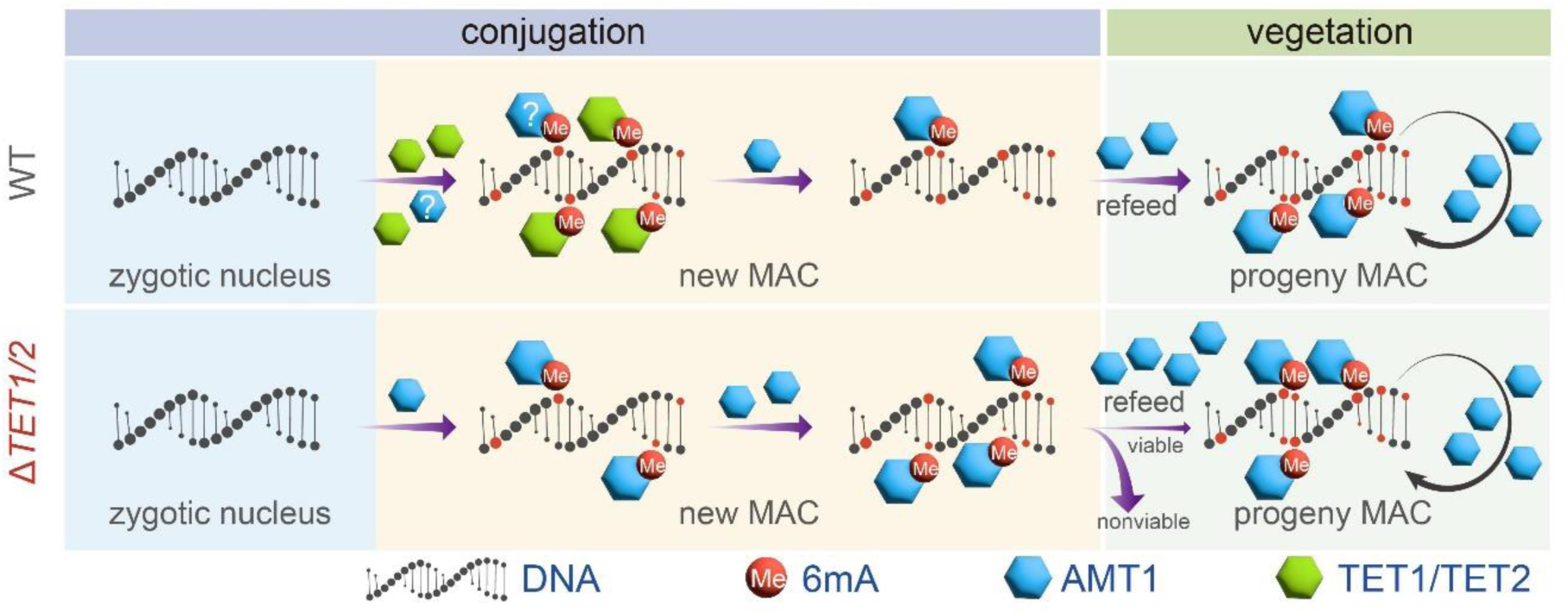
Proposed model for 6mA establishment and maintenance in WT and KO cells. In the new MAC of WT cells, TET1 and TET2 initiated the *de novo* 6mA deposition on unmethylated ApT dinucleotides, possibly helped by AMT1, generating hemi-6mApT. Subsequently, these hemi-6mApT sites were converted into full-6mApT by the maintenance activity of AMT1. In the somatic MAC during the vegetative stage, established full-6mApT sites were faithfully maintained by AMT1. In *TET1* and/or *TET2* KO cells, however, the deposition of *de novo* 6mA dependent on TET1 and TET2 was abolished, while AMT1 could exhibit both *de novo* and maintenance activities and generate both hemi- and full-6mA on a limited number of sites. After refeeding, 6mA levels in KO cells could be restored to that of WT cells, probably owing to the upregulated level of AMT1.

In *TET1* and/or *TET2* KO cells, the occurrence of 6mA was not recovered until refeeding. During the late stage of conjugation without refeeding, AMT1 activity likely remained low, modulated by both its low protein level and the possible absence or low levels of its interacting proteins. After refeeding, however, AMT1 level was dramatically upregulated, probably for fulfilling its role as a maintenance MTase ^29^. The elevated level of AMT1 successfully restored global 6mA levels in KO cells to that of WT cells, although we could not rule out the possibility that a small proportion of sites that relied solely on TET1/TET2 might remain unmethylated. Given that very few (endo)replication was observed during late conjugation (Figure S8C) ^42,43^, in contrast to active replication during the vegetative stage ^44^, we speculated that AMT1 level and/or activity might also be linked with replication.

Together, our work identified and characterized the primary *de novo* MTase for eukaryotic 6mA, affirming the longstanding hypothesis that 6mA dynamics also involve a two-step pathway. These findings, combined with our previous discoveries regarding AMT1, delineate a detailed pathway for the proper establishment and maintenance of 6mA.

### Comparing 6mA and 5mC

There are several key similarities between *de novo* 6mA in the new MAC of *Tetrahymena* and *de novo* 5mC in the early embryogenesis of mammalian cells. First, both modifications occur during the transition from a zygotic nucleus to a somatic nucleus ^11,52,53^, accomplished by intensive epigenome reprogramming ^54–56^. Second, in both cases, *de novo* DNA methylation is catalyzed by specific DNA MTase(s), TET1 and TET2 in the case of 6mA and DNMT3A and DNMT3B for 5mC ^19,57^. Third, the *de novo* depositions of both 6mA and 5mC reply on a complex interplay with other epigenetic factors to establish specific patterns. For 6mA, active transcription marks such as H2A.Z and H3K4me3, and possibly histone acetylation, are the major players ^11,28^. Repressive marks such as H3K9me3 and H3K27me3 might also play a role to confine the territory of 6mA ^58,59^. For 5mC, the concerto playing with H3K9me2/me3, as well as histone deacetylase, confers its genome targeting and reinforces its silencing effects ^60–62^. Fourth, both modifications play essential roles for proper development. Loss of TET1 and/or TET2 reduces the progeny viability in *Tetrahymena*, while the absence of DNMT3A and DNMT3B results in aberrant gene expression and embryo lethality in mammals ^19,63–65^.

It is also important to note that while there are shared features, the distinctions between 6mA in *Tetrahymena* and 5mC in mammals are significant, arising from their unique evolutionary background and biological contexts. First, the *Tetrahymena* zygotic nucleus originates from the 6mA-free micronucleus ^30–32^, thus requiring no demethylation prior to *de novo* 6mA deposition. Conversely, genome-wide *de novo* 5mC deposition occur in parallel or after the germline 5mC was erased by both passive and active demethylation ^66–68^. Second, the *de novo* MTases for 6mA and 5mC have distinct enzymatic features. TET1 and TET2 convert unmethylated adenines to hemi-6mA, while DNMT3A and DNMT3B generate both hemi- and full-5mC ^69,70^. Third, in a fully methylated *Tetrahymena* somatic genome, only ∼2% of ApT are methylated ^11,29^, while 60-80% CpG are methylated in mammalian somatic genomes ^71^. Fourth, 6mA is closely associated with active transcription, fulfilling its role, together with other epigenetic factors such as H2A.Z and H3K4me3 ^11,13,28^ to activate the somatic transcription, so that *Tetrahymena* could thrive as a single cell executing a variety of functions such as feeding, mating, etc. In contrast, 5mC is primarily linked with gene repression, preventing the expression of genes unnecessary in specific cell types, besides limiting transposable elements and repetitive sequences ^72–75^.

In summary, our work established a notable parallel between 6mA and 5mC: both feature palindrome motifs (ApT vs. CpG), transmit in a semi-conservative manner, and require a two-step methylation pathway. Considering the dichotomy of 6mA prevalence in unicellular eukaryotes versus 5mC in multicellular eukaryotes, our study offers new insights into the evolutionary divergence among eukaryotes from the unique perspective of DNA methylation.

## METHODS AND MATERIALS

### Cell culture

*Tetrahymena thermophila* wild-type strains, SB210 and CU428, were obtained from the *Tetrahymena* Stock Center (http://tetrahymena.vet.cornell.edu). *AMT1* germline knockout (KO) homozygous homokaryon strain was generated in our previous work ^11^. KO and tag strains used in this study, most of which were newly generated, were listed in Table S5. Cells were grown in SPP medium at _30°C_ ^76,77^.

### Generation of *Tetrahymena* strains

Primers used in strain generation and confirmation were listed in Table S6.

To generate KO constructs, *neo4* or *bsr2* cassettes ^78,79^ were flanked with 5’ and 3’ flanking regions of *TET1* and *TET2* (Figure S1A). *TET1* and *TET2* germline single or double KO homozygous homokaryons and homozygous heterokaryons (Δ*TET1* and Δ*TET2*) were generated according to reported procedures (Figures S1B-C)^11^. Somatic KO strains were generated as previously reported (Figure S1B) ^80,81^.

To generate tagging constructs, sequences coding for the hemagglutinin (HA) tag were inserted to the C-terminus of the *TET1* or the N-terminus of *TET2* and *AMT1* (Figure S3A). Germline tagging strain of TET1 (TET1-CHA) and somatic tagging strain of TET2 (TET2-NHA) were generated as previously reported ^82^ (Figure S3B). Germline tagging strain of AMT1-NHA in the background of WT cells (SB210 and CU428) was previously generated ^83^, while those in the background of *TET1* and/or *TET2* KO cells were generated by crossing two different mating types of homozygous heterokaryon AMT1-NHA with homozygous heterokaryons deleting each or both of *TET1* and *TET2* following the same strategy used for generating double KO cells (Figure S1B).

To generate *TET1*-rescue (*TET1-*RS) and *TET2*-rescue (*TET2*-RS) constructs, the full-length *TET1* or *TET2* ORF and its flanking regions were cloned and the *bsr2* cassette was inserted into the 3’ UTR. The rescue construct was transformed into the MAC of corresponding germline KO cells and selected by blasticidin S (Figure S4).

To generate TET1 mutants, a sequence spanning the 5’ flanking region, the full-length TET1 with HA tag, the 3’ flanking region, and the *bsr2* cassette was PCR amplified from the somatic *TET1*-RS cells. The mutation was introduced by fusion PCR with primers containing the designed mutations (Table S6). The mutant construct was transformed into the somatic MAC of Δ*TET1* cells and selected by blasticidin S (Figure S4).

To distinguish true progeny from quitter, the *RPB3*-CHA-bsr2 construct with a HA tag at the C-terminus of *RPB3* was transformed into the somatic MAC of WT, Δ*TET1*, Δ*TET2*, and Δ*TET1/2* cells (Figures 6E and S5A).

### Immunofluorescence staining

Different fixation methods using 2% paraformaldehyde (Sigma, P6148), combination of saturated HgCl_2_ (HQ, HG 3-1068-77) and ethanol (1:2), or 10% formalin (Nacalai Tesque, 16223-55) were used as previously described ^11,28,41,80,84–87^. Conjugative cells were fixed at indicated timepoints after mixing cells of two different mating types. The primary antibodies were ɑ-6mA (Synaptic Systems, 202003, 1:2000, rabbit) and ɑ-HA (Cell Signaling, C29F4, 1:200, rabbit). The secondary antibodies were Goat anti-Rabbit IgG (H+L), Alexa Fluor 555 (Invitrogen, A-21428, 1:4000), Goat anti-Rabbit IgG (H+L), Alexa Fluor 546 (Invitrogen, A-11035, 1:4000), and Goat anti-Mouse IgG (H+L), Alexa Fluor 488 (Invitrogen, A-11001, 1:4000).

### Purification of new MACs

The new MACs were isolated from conjugative WT and KO cells by differential centrifugation following previous procedures ^88^. Briefly, nuclear pellets containing a large proportion of new MACs were collected at 3,500-4,500g from 24h post-mixing cells. Nuclear pellets were washed once with 4′,6-Diamidine-2′-phenylindole dihydrochloride (DAPI) (Roche, 10236276001) in methanol (1 μg/ml), then resuspended in DAPI-methanol (1 μg/ml) and incubated in the dark at room temperature for 15 minutes. Nuclei were resuspended in modified TMSN buffer (0.25 M sucrose, 10 mM Tris-HCl pH 7.5, 3 mM CaCl_2_, 0.016% NP-40, 1× complete proteinase inhibitor cocktail (Roche, 11873580001)) ^41^ and sorted by Becton Dickinson FACS Aria III flow cytometer (Becton, Dickinson and Company, Pleasanton, CA, United States). The purity of the collected new MACs was assessed by DAPI-stained nuclear morphology. MICs were easily distinguished by their much smaller size; somatic MACs and new MACs were distinguishable by their differences in size (bigger vs. smaller) and DAPI signal intensity (stronger vs. weaker). Contamination rate of somatic MACs and MICs in the new MAC fraction were < 2% and < 10%, respectively.

### UHPLC–QQQ–MS/MS analysis

Genomic DNA was extracted using DNA Extraction Reagent (Solarbio, P1012) or QIAGEN Genomic-tip 20/G (10223) and then processed into single nucleotides as previously described ^11^. These single nucleotides were analyzed by ultra-high-performance liquid chromatography tandem mass spectrometry (UHPLC–QQQ–MS/MS) on an Acquity BEH C18 column (100 mm × 2.1 mm, 1.7 µm, Waters, MA, USA) using a Xevo TQ-S triple quadrupole mass spectrometer (Waters, Milford, MA, USA), or a Hypersil GOLD column (100 × 2.1 mm, 1.9 µm, Thermo scientific) using a TSQ Quantiva mass spectrometer (Thermo scientific). The mass spectrometer was set to multiple reaction monitoring (MRM) under positive electrospray mode. For 6mA and dA, the selective MRM transitions were detected under m/z 266/150 and m/z 252/136, respectively. The ratio of 6mA/A was quantified by the running nucleoside standards calibration curves at the same time.

### Conjugation progress analysis and viability test

To monitor the conjugation progress, cells at different stages (4h, 6h, 10h, 14h, 24h after mixing) of Δ*TET1* (mating types IV and V), Δ*TET2* (mating types II and IV), Δ*TET1/*Δ*TET2* (mating types II and IV), and WT (SB210 and CU428) were fixed with 2% paraformaldehyde, stained with 1 μg/mL DAPI (BBI, E607303), and observed under the microscope.

For the viability test of each KO strain (WT cells were used as control), individual mating pair was picked into SPP medium drops at 10h post-mixing. After 2∼3 days culture, cells were transferred into 96-well plates. It should be noted that the blasticidin S resistance was introduced into the parental MAC of mating cells by transforming a *bsr2* cassette-containing construct ^79^, which could help to distinguish true progenies (blasticidin S sensitive) from aborted pairs (quitters, blasticidin S resistant) (Figure S5A). Viable cells in 96-well plates that were sensitive to blasticidin S were counted as viable true progenies. True progenies possessed mating types other than that of parental cells, demonstrated by PCR amplification-based test using mating type primers (Figure S5B, Table S6) ^89^, confirming the efficacy of blasticidin S selection.

### SMRT sequencing and data analysis

Genomic DNA was extracted from FACS-sorted new MACs of both WT and KO cells using DNA Extraction Reagent (Solarbio, P1012). The quality and concentration of DNA were assessed by agarose gel electrophoresis and the Qubit^®^3.0 Fluorometer (Thermo Fisher Scientific).

Sample preparation and data analysis followed the latest CCS analysis pipeline ^29^. Genomic DNA was sheared to approximately 3kb to generate PacBio Sequel II libraries, ensuring sufficient subreads coverage and sequencing depth. To evaluate the sequencing quality, samtools ^90^ was employed to sort the subreads and the ccs module (SMRT Link v11.0, Pacific Biosciences) was used to generate circular consensus sequence and sequencing information for each single molecule. Customed Perl scripts were then used to extract high-confidence single molecules (passes ≥ 30×) for downstream analyses.

Single molecules were mapped to their corresponding consensus sequence using BLASR ^91^ and IPD ratios were calculated using ipdSummary (SMRT Link v11.0, Pacific Biosciences). Single molecules with global dispersion, defined as large IPD ratios standard deviation (SD) in Watson and/or Crick strands (SD ≥ 0.35) for all unmethylated adenine sites (IPD ratios < 2.8), were filtered out. To minimize the impact of sequencing variability, single molecules with local dispersion of IPD ratios, characterized by high-density of N*, were filtered out ^29^.

Circular consensus sequences of selected single molecules were mapped back to the *Tetrahymena* genome assembly, which contained sequences from the MIC, MAC, and mitochondrion genomes ^92–95^, using blastn ^96^ with the parameter “-max_hsps 1, -max_target_seqs 1”. MAC-mapped reads (mapped length ≥ 98% for read length, mapped identity ≥ 95%) were selected for further analysis. Potential MIC reads contaminations were detected based on the alignment score difference between MIC and MAC (Δ ≥ 50) ^29^. 6mA IPD ratio threshold of all single molecules were calculated by applying the bimodal distribution deconvolution to IPD ratios.

### RNA sequencing and data analysis

Progenies of Δ*TET1* (mating type IV and V), Δ*TET2* (mating type II and IV), Δ*TET1/2* (mating type II and IV) and WT (SB210 and CU428) cells were distinguished from non-maters and quitters based on their resistance to blasticidin S (Figure S5A). Total RNA from two replicates of each strain were extracted at about 20 generations after refeeding using RNAiso Plus (TaKaRa, 9108).

After trimming sequencing adapters and filtering out low quality reads by Trim Galore ^97^, the resulting pair-end reads were mapped to the latest MAC genome assembly in the *Tetrahymena* genome database (TGD) (http://ciliate.org) ^92,95^ using HISAT2 ^98^. Discordant alignments for paired reads were discarded. Only paired reads with unique mapping, as indicated by the presence of the ‘NH:i:1’ tag in the mapped SAM file, were retained. PCR duplicates were removed using the Picard MarkDuplicates tool (http://broadinstitute.github.io/picard/). The featureCounts program ^99^ was used for counting reads mapped to genomic features, using gene annotation file from TGD as the reference. The average reads for each gene in every strain were calculated to elucidate the similarity of transcription profiles. The Spearman’s correlation coefficient between every two strains was calculated using R, and the heatmap was generated using the gplots package ^100^.

### Data availability

The latest SB210 MAC genome can be found at the *Tetrahymena* genome database (TGD) (http://ciliate.org) ^92,95^. RNA-seq data used in this study were deposited at the NCBI database (PRJNA1071335).

## ACKNOWLEDGEMENTS

This work is supported by the National Natural Science Foundation of China (32125006, 32070437) and the Science & Technology Innovation Project of Laoshan Laboratory (LSKJ202203203). The authors would like to thank the following people for assistance with this study: Dr. Kai Liu (College of Life Sciences, Wuhan University, China) for suggestions on protein purification, Dr. Zhiyi Lv for fruitful discussion, and Ms. Jiaoxia Shi and Ms. Yuanyuan Zhang (Ocean University of China (OUC), China) for assisting the flow cytometry sorting. High-performance computing resources for data processing were provided by the Institute of Evolution & Marine Biodiversity at OUC, the Center for High Performance Computing and System Simulation at Laoshan Laboratory, and Marine Big Data Center of Institute for Advanced Ocean Study at OUC. Our special thanks are given to Prof. Weibo Song (OUC) for his helpful suggestions during drafting the manuscript.

## AUTHOR CONTRIBUTIONS

T.C. conceived and led the project, performed most of the experiments, prepared figures, and wrote and revised the manuscript. J.Z. and J.D. performed the experiments, prepared figures, and revised the manuscript. H.L. performed the bioinformatic analysis, prepared figures, and revised the manuscript. W.Z. and J.N. performed the mass spectrometry analysis. K.K. supervised the generation and characterization of several mutant strains and revised the manuscript. S.G. conceived and supervised the project, wrote and revised the manuscript, and provided funding resources. All authors read and approved the final manuscript.

## REFERENCES

1. Fu, Y., Luo, G., Chen, K., Deng, X., Yu, M., Han, D., Hao, Z., Liu, J., Lu, X., Dore, L.C., et al. (2015). N^6^-methyldeoxyadenosine marks active transcription start sites in *Chlamydomonas*. Cell 161, 879–892.

2. Greer, E.L., Blanco, M.A., Gu, L., Sendinc, E., Liu, J., Aristizábal-Corrales, D., Hsu, C., Aravind, L., He, C., and Shi, Y. (2015). DNA methylation on N^6^-adenine in *C. elegans*. Cell 161, 868–878.

3. Zhang, G., Huang, H., Liu, D., Cheng, Y., Liu, X., Zhang, W., Yin, R., Zhang, D., Zhang, P., Liu, J., et al. (2015). N^6^-methyladenine DNA modification in *Drosophila*. Cell 161, 893–906.

4. Hao, Z., Wu, T., Cui, X., Zhu, P., Tan, C., Dou, X., Hsu, K., Lin, Y., Peng, P., Zhang, L., et al. (2020). N^6^-Deoxyadenosine methylation in mammalian mitochondrial DNA. Mol. Cell 78, 382–395.e388.

5. Zhang, M., Yang, S., Nelakanti, R., Zhao, W., Liu, G., Li, Z., Liu, X., Wu, T., Xiao, A., and Li, H. (2020). Mammalian ALKBH1 serves as an N^6^-mA demethylase of unpairing DNA. Cell Res. 30, 197–210.

6. He, S., Zhang, G., Wang, J., Gao, Y., Sun, R., Cao, Z., Chen, Z., Zheng, X., Yuan, J., Luo, Y., et al. (2019). 6mA-DNA-binding factor Jumu controls maternal-to-zygotic transition upstream of Zelda. Nat. Commun. 10, 2219.

7. Liu, Y., Niu, J., Ye, F., Solberg, T., Lu, B., Wang, C., Nowacki, M., and Gao, S. (2024). Dynamic DNA *N*^6^-adenine methylation (6mA) governs the encystment process, showcased in the unicellular eukaryote *Pseudocohnilembus persalinus*. Genome Res.

8. Wu, T.P., Wang, T., Seetin, M.G., Lai, Y., Zhu, S., Lin, K., Liu, Y., Byrum, S.D., Mackintosh, S.G., Zhong, M., et al. (2016). DNA methylation on N^6^-adenine in mammalian embryonic stem cells. Nature 532, 329–333.

9. Zhang, S., Li, B., Du, K., Liang, T., Dai, M., Huang, W., Zhang, H., Ling, Y., and Zhang, H. (2020). Epigenetically modified N^6^-methyladenine inhibits DNA replication by human DNA polymerase iota. Biochimie 168, 134–143.

10. Zhang, X., Blumenthal, R.M., and Cheng, X. (2021). A role for N^6^-methyladenine in DNA damage repair. Trends Biochem. Sci. 46, 175–183.

11. Wang, Y., Sheng, Y., Liu, Y., Zhang, W., Cheng, T., Duan, L., Pan, B., Qiao, Y., Liu, Y., and Gao, S. (2019). A distinct class of eukaryotic MT-A70 methyltransferases maintain symmetric DNA N^6^-adenine methylation at the ApT dinucleotides as an epigenetic mark associated with transcription. Nucleic Acids Res. 47, 11771–11789.

12. Beh, L.Y., Debelouchina, G.T., Clay, D.M., Thompson, R.E., Lindblad, K.A., Hutton, E.R., Bracht, J.R., Sebra, R.P., Muir, T.W., and Landweber, L.F. (2019). Identification of a DNA N^6^-adenine methyltransferase complex and its impact on chromatin organization. Cell 177, 1781–1796.e1725.

13. Ma, C., Niu, R., Huang, T., Shao, L.W., Peng, Y., Ding, W., Wang, Y., Jia, G., He, C., Li, C.Y., et al. (2019). N^6^-methyldeoxyadenine is a transgenerational epigenetic signal for mitochondrial stress adaptation. Nat. Cell Biol. 21, 319–327.

14. Otto, G. (2019). 6mA in stress tolerance across generations. Nat. Rev. Mol. Cell Biol. 20, 66–67.

15. Yao, B., Cheng, Y., Wang, Z., Li, Y., Chen, L., Huang, L., Zhang, W., Chen, D., Wu, H., Tang, B., and Jin, P. (2017). DNA N^6^-methyladenine is dynamically regulated in the mouse brain following environmental stress. Nat. Commun. 8, 1122.

16. Liu, J., Zhu, Y., Luo, G., Wang, X., Yue, Y., Wang, X., Zong, X., Chen, K., Yin, H., Fu, Y., et al. (2016). Abundant DNA 6mA methylation during early embryogenesis of zebrafish and pig. Nat. Commun. 7, 13052.

17. Liang, Z., Shen, L., Cui, X., Bao, S., Geng, Y., Yu, G., Liang, F., Xie, S., Lu, T., Gu, X., and Yu, H. (2018). DNA N^6^-Adenine methylation in *Arabidopsis thaliana*. Dev. Cell 45, 406–416.e403.

18. Xie, Q., Wu, T.P., Gimple, R.C., Li, Z., Prager, B.C., Wu, Q., Yu, Y., Wang, P., Wang, Y., Gorkin, D.U., et al. (2018). N^6^-methyladenine DNA modification in glioblastoma. Cell 175, 1228–1243.e1220.

19. Okano, M., Bell, D., Haber, D., and Li, E. (1999). DNA methyltransferases Dnmt3a and Dnmt3b are essential for *de novo* methylation and mammalian development. Cell 99, 247–257.

20. Yoder, J.A., Soman, N.S., Verdine, G.L., and Bestor, T.H. (1997). DNA (cytosine-5)-methyltransferases in mouse cells and tissues. Studies with a mechanism-based probe. J. Mol. Biol. 270, 385–395.

21. Stein, R., Gruenbaum, Y., Pollack, Y., Razin, A., and Cedar, H. (1982). Clonal inheritance of the pattern of DNA methylation in mouse cells. Proc. Natl. Acad. Sci. USA 79, 61–65.

22. Howlett, S.K., and Reik, W. (1991). Methylation levels of maternal and paternal genomes during preimplantation development. Development 113, 119–127.

23. Kafri, T., Ariel, M., Brandeis, M., Shemer, R., Urven, L., Mccarrey, J., Cedar, H., and Razin, A. (1992). Developmental pattern of gene-specific DNA methylation in the mouse embryo and germ line. Genes Dev. 6, 705–714.

24. Monk, M., Boubelik, M., and Lehnert, S. (1987). Temporal and regional changes in DNA methylation in the embryonic, extraembryonic and germ cell lineages during mouse embryo development. Development 99, 371–382.

25. Sanford, J.P., Clark, H.J., Chapman, V.M., and Rossant, J. (1987). Differences in DNA methylation during oogenesis and spermatogenesis and their persistence during early embryogenesis in the mouse. Genes Dev. 1, 1039–1046.

26. Chen, T., Ueda, Y., Dodge, J.E., Wang, Z., and Li, E. (2003). Establishment and maintenance of genomic methylation patterns in mouse embryonic stem cells by Dnmt3a and Dnmt3b. Mol. Cell. Biol. 23, 5594–5605.

27. Li, E., Bestor, T.H., and Jaenisch, R. (1992). Targeted mutation of the DNA methyltransferase gene results in embryonic lethality. Cell 69, 915–926.

28. Wang, Y., Chen, X., Sheng, Y., Liu, Y., and Gao, S. (2017). N^6^-adenine DNA methylation is associated with the linker DNA of H2A.Z-containing well-positioned nucleosomes in Pol II-transcribed genes in *Tetrahymena*. Nucleic Acids Res. 45, 11594–11606.

29. Sheng, Y., Wang, Y., Yang, W., Wang, X.Q., Lu, J., Pan, B., Nan, B., Liu, Y., Li, C., Song, J., et al. (2023). Semi-conservative transmission of DNA N^6^-adenine methylation in a unicellular eukaryote. bioRxiv, 2023.2002.2015.468708.

30. Gorovsky, M.A., and Pleger, H. (1973). [6N]methyl adenine in the nuclear DNA of a eucaryote, *Tetrahymena pyriformis*. J. Mol. Biol. 56, 697–701.

31. Harrison, G.S., and Karrer, K.M. (1985). DNA synthesis, methylation and degradation during conjugation in *Tetrahymena thermophila*. Nucleic Acids Res. 13, 73–87.

32. Harrison, G.S., Findly, R.C., and Karrer, K.M. (1986). Site-specific methylation of adenine in the nuclear genome of a eucaryote, *Tetrahymena thermophila*. Mol. Cell. Biol. 6, 2364–2370.

33. Karrer, K.M., and Vannuland, T.A. (2002). Methylation of adenine in the nuclear DNA of *Tetrahymena* is internucleosomal and independent of histone H1. Nucleic Acids Res. 30, 1364–1370.

34. Wei, F., Pan, B., Diao, J., Wang, Y., Sheng, Y., and Gao, S. (2022). The micronuclear histone H3 clipping in the unicellular eukaryote *Tetrahymena thermophila*. Mar. Life Sci. Technol. 4, 584–594.

35. Tian, M., Cai, X., Liu, Y., Liucong, M., and Howard-Till, R. (2022). A practical reference for studying meiosis in the model ciliate *Tetrahymena thermophila*. Mar. Life Sci. Technol. 4, 595–608.

36. Zhao, X., Li, Y., Duan, L., Chen, X., Mao, F., Juma, M., Liu, Y., Song, W., and Gao, S. (2020). Functional analysis of the methyltransferase SMYD in the single-cell model organism *Tetrahymena thermophila*. Mar. Life Sci. Technol. 2, 109–122.

37. Mondo, S.J., Dannebaum, R.O., Kuo, R.C., Louie, K.B., Bewick, A.J., LaButti, K., Haridas, S., Kuo, A., Salamov, A., Ahrendt, S.R., et al. (2017). Widespread adenine N^6^-methylation of active genes in fungi. Nat. Genet. 49, 964–968.

38. Bochtler, M., and Fernandes, H. (2021). DNA adenine methylation in eukaryotes: Enzymatic mark or a form of DNA damage? Bioessays 43, e2000243.

39. Miao, W., Xiong, J., Bowen, J., Wang, W., Liu, Y., Braguinets, O., Grigull, J., Pearlman, R.E., Orias, E., and Gorovsky, M.A. (2009). Microarray analyses of gene expression during the *Tetrahymena thermophila* life cycle. PLoS One 4, e4429.

40. Stargell, L.A., Bowen, J., Dadd, C.A., Dedon, P.C., Davis, M., Cook, R.G., Allis, C.D., and Gorovsky, M.A. (1993). Temporal and spatial association of histone H2A variant hv1 with transcriptionally competent chromatin during nuclear development in *Tetrahymena thermophila*. Genes Dev. 7, 2641–2651.

41. Kataoka, K., and Mochizuki, K. (2015). Phosphorylation of an HP1-like protein regulates heterochromatin body assembly for DNA elimination. Dev. Cell 35, 775–788.

42. Allis, C.D., Colavito-Shepanski, M., and Gorovsky, M.A. (1987). Scheduled and unscheduled DNA synthesis during development in conjugating *Tetrahymena*. Dev. Biol. 124, 469–480.

43. Yin, L., Gater, S.T., and Karrer, K.M. (2010). A developmentally regulated gene, ASI2, is required for endocycling in the macronuclear anlagen of *Tetrahymena*. Eukaryot. Cell 9, 1343–1353.

44. McDonald, B.B. (1962). Synthesis of deoxyribonucleic acid by micro- and macronuclei of *Tetrahymena pyriformis*. J. Cell Biol. 13, 193–203.

45. 45. Lynn, D.H. (2008). The Ciliated Protozoa: Characterization, Classification, and Guide to the Literature, Third Edition (Springer Science and Business Media B.V.).

46. Xiong, J., Yang, W., Chen, K., Jiang, C., and Miao, W. (2019). Hidden genomic evolution in a morphospecies—The landscape of rapidly evolving genes in *Tetrahymena*. PLoS Biol. 17, e3000294.

47. Chen, J., Hu, R., Chen, Y., Lin, X., Xiang, W., Chen, H., Yao, C., and Liu, L. (2022). Structural basis for MTA1c-mediated DNA N^6^-adenine methylation. Nat. Commun. 13, 3257.

48. Woodcock, C.B., Horton, J.R., Zhang, X., Blumenthal, R.M., and Cheng, X. (2020). Beta class amino methyltransferases from bacteria to humans: evolution and structural consequences. Nucleic Acids Res. 48, 10034–10044.

49. Yan, J., Liu, F., Guan, Z., Yan, X., Jin, X., Wang, Q., Wang, Z., Yan, J., Zhang, D., and Liu, Z. (2023). Structural insights into DNA N^6^-adenine methylation by the MTA1 complex. Cell Discov. 9, 8.

50. Liu, J., Yue, Y., Han, D., Wang, X., Fu, Y., Zhang, L., Jia, G., Yu, M., Lu, Z., and Deng, X. (2014). A METTL3–METTL14 complex mediates mammalian nuclear RNA N^6^-adenosine methylation. Nat. Chem. Biol. 10, 93–95.

51. Wang, P., Doxtader, K.A., and Nam, Y. (2016). Structural basis for cooperative function of Mettl3 and Mettl14 methyltransferases. Mol. Cell 63, 306–317.

52. Li, L., Lu, X., and Dean, J. (2013). The maternal to zygotic transition in mammals. Mol. Aspects Med. 34, 919–938.

53. Martindale, D.W., Allis, C.D., and Bruns, P.J. (1982). Conjugation in *Tetrahymena thermophila*: a temporal analysis of cytological stages. Exp. Cell Res. 140, 227–236.

54. Reik, W., Dean, W., and Walter, J. (2001). Epigenetic reprogramming in mammalian development. Science 293, 1089–1093.

55. Meyer, E., and Duharcourt, S. (1996). Epigenetic programming of developmental genome rearrangements in ciliates. Cell 87, 9–12.

56. Nowacki, M., Shetty, K., and Landweber, L.F. (2011). RNA-mediated epigenetic programming of genome rearrangements. Annu. Rev. Genomics Hum. Genet. 12, 367–389.

57. Hsieh, C.L. (1999). *In vivo* activity of murine *de novo* methyltransferases, Dnmt3a and Dnmt3b. Mol. Cell. Biol. 19, 8211–8218.

58. Noto, T., and Mochizuki, K. (2018). Small RNA-mediated *trans*-nuclear and *trans*-element communications in *Tetrahymena* DNA elimination. Curr. Biol. 28, 1938–1949.e1935.

59. Suhren, J.H., Noto, T., Kataoka, K., Gao, S., Liu, Y., and Mochizuki, K. (2017). Negative regulators of an RNAi-heterochromatin positive feedback loop safeguard somatic genome integrity in *Tetrahymena*. Cell Rep. 18, 2494–2507.

60. Lehnertz, B., Ueda, Y., Derijck, A.A., Braunschweig, U., Perez-Burgos, L., Kubicek, S., Chen, T., Li, E., Jenuwein, T., and Peters, A.H. (2003). Suv39h-mediated histone H3 lysine 9 methylation directs DNA methylation to major satellite repeats at pericentric heterochromatin. Curr. Biol. 13, 1192–1200.

61. 61. Fouse, S.D., Shen, Y., Pellegrini, M., Cole, S., Meissner, A., Van Neste, L., Jaenisch, R., and Fan, G. (2008). Promoter CpG methylation contributes to ES cell gene regulation in parallel with Oct4/Nanog, PcG complex, and histone H3 K4/K27 trimethylation. Cell stem cell 2, 160–169.

62. Fuks, F., Hurd, P.J., Deplus, R., and Kouzarides, T. (2003). The DNA methyltransferases associate with HP1 and the SUV39H1 histone methyltransferase. Nucleic Acids Res. 31, 2305–2312.

63. Dodge, J.E., Okano, M., Dick, F., Tsujimoto, N., Chen, T., Wang, S., Ueda, Y., Dyson, N., and Li, E. (2005). Inactivation of Dnmt3b in mouse embryonic fibroblasts results in DNA hypomethylation, chromosomal instability, and spontaneous immortalization. J. Biol. Chem. 280, 17986–17991.

64. Kaneda, M., Okano, M., Hata, K., Sado, T., Tsujimoto, N., Li, E., and Sasaki, H. (2004). Essential role for *de novo* DNA methyltransferase Dnmt3a in paternal and maternal imprinting. Nature 429, 900–903.

65. 65. Rinaldi, L., Datta, D., Serrat, J., Morey, L., Solanas, G., Avgustinova, A., Blanco, E., Pons, J.I., Matallanas, D., Von Kriegsheim, A., et al. (2016). Dnmt3a and Dnmt3b associate with enhancers to regulate human epidermal stem cell homeostasis. Cell stem cell 19, 491–501.

66. Kohli, R.M., and Zhang, Y. (2013). TET enzymes, TDG and the dynamics of DNA demethylation. Nature 502, 472–479.

67. Wu, X., and Zhang, Y. (2017). TET-mediated active DNA demethylation: mechanism, function and beyond. Nat. Rev. Genet. 18, 517–534.

68. Gong, Z., and Zhu, J.K. (2011). Active DNA demethylation by oxidation and repair. Cell Res. 21, 1649–1651.

69. Li, Z., Dai, H., Martos, S.N., Xu, B., Gao, Y., Li, T., Zhu, G., Schones, D.E., and Wang, Z. (2015). Distinct roles of DNMT1-dependent and DNMT1-independent methylation patterns in the genome of mouse embryonic stem cells. Genome Biol. 16, 115.

70. Hsieh, C.L. (2005). The *de novo* methylation activity of Dnmt3a is distinctly different than that of Dnmt1. BMC Biochem. 6, 1–12.

71. Smith, Z.D., and Meissner, A. (2013). DNA methylation: roles in mammalian development. Nat. Rev. Genet. 14, 204–220.

72. Pastor, W.A., Stroud, H., Nee, K., Liu, W., Pezic, D., Manakov, S., Lee, S.A., Moissiard, G., Zamudio, N., Bourc’his, D., et al. (2014). MORC1 represses transposable elements in the mouse male germline. Nat. Commun. 5, 5795.

73. Rosic, S., Amouroux, R., Requena, C.E., Gomes, A., Emperle, M., Beltran, T., Rane, J.K., Linnett, S., Selkirk, M.E., Schiffer, P.H., et al. (2018). Evolutionary analysis indicates that DNA alkylation damage is a byproduct of cytosine DNA methyltransferase activity. Nat. Genet. 50, 452–459.

74. Deniz, Ö., Frost, J.M., and Branco, M.R. (2019). Regulation of transposable elements by DNA modifications. Nat. Rev. Genet. 20, 417–431.

75. Kim, M., Park, Y.K., Kang, T.W., Lee, S.H., Rhee, Y.H., Park, J.L., Kim, H.J., Lee, D., Lee, D., Kim, S.Y., and Kim, Y.S. (2014). Dynamic changes in DNA methylation and hydroxymethylation when hES cells undergo differentiation toward a neuronal lineage. Hum. Mol. Genet. 23, 657–667.

76. Gorovsky, M.A., Yao, M.C., Keevert, J.B., and Pleger, G.L. (1975). Isolation of micro- and macronuclei of *Tetrahymena pyriformis*. Methods Cell Biol. 9, 311–327.

77. Sweet, M., and Allis, C.D. (1998). Cells: A Laboratory Manual. Spector, DL, Goldman, RD and Leinwand, LA (eds), Volume 1 (Cold Spring Harbor Laboratory Press).

78. Mochizuki, K. (2008). High efficiency transformation of *Tetrahymena* using a codon-optimized neomycin resistance gene. Gene 425, 79–83.

79. Talsky, K.B., and Collins, K. (2012). Strand-asymmetric endogenous *Tetrahymena* small RNA production requires a previously uncharacterized uridylyltransferase protein partner. RNA 18, 1553–1562.

80. Gao, S., Xiong, J., Zhang, C., Berquist, B.R., Yang, R., Zhao, M., Molascon, A.J., Kwiatkowsk, S.Y., Yuan, D., Qin, Z., et al. (2013). Impaired replication elongation in *Tetrahymena* mutants deficient in histone H3 Lys 27 monomethylation. Genes Dev. 27, 1662–1679.

81. Qiao, Y., Cheng, T., Zhang, J., Alfarraj, S.A., Tian, M., Liu, Y., and Gao, S. (2022). Identification and utilization of a mutated 60S ribosomal subunit coding gene as an effective and cost-efficient selection marker for *Tetrahymena* genetic manipulation. Int. J. Biol. Macromol. 204, 1–8.

82. Yu, L., and Gorovsky, M.A. (1999). Protein tagging in *Tetrahymena*. Methods Cell Biol. 62, 549–559.

83. Duan, L., Li, H., Ju, A., Zhang, Z., Niu, J., Zhang, Y., Diao, J., Kataoka, K., Ma, H., Song, N., et al. (2024). Self-regulation of the DNA N^6^-adenine methyltransferase AMT1 in the unicellular eukaryote *Tetrahymena thermophila*. bioRxiv, 2024.2002.2006.579081.

84. Xu, J., Li, X., Song, W., Wang, W., and Gao, S. (2019). Cyclin Cyc2p is required for micronuclear bouquet formation in *Tetrahymena thermophila*. Sci. China Life Sci. 62, 668–680.

85. Zhao, X., Wang, Y., Wang, Y., Liu, Y., and Gao, S. (2017). Histone methyltransferase TXR1 is required for both H3 and H3. 3 lysine 27 methylation in the well-known ciliated protist *Tetrahymena thermophila*. Sci. China Life Sci. 60, 264–270.

86. Loidl, J., and Mochizuki, K. (2009). *Tetrahymena* meiotic nuclear reorganization is induced by a checkpoint kinase-dependent response to DNA damage. Mol. Biol. Cell 20, 2428–2437.

87. Loidl, J., and Scherthan, H. (2004). Organization and pairing of meiotic chromosomes in the ciliate *Tetrahymena thermophila*. J. Cell Sci. 117, 5791–5801.

88. Feng, L., Wang, G., Hamilton, E.P., Xiong, J., Yan, G., Chen, K., Chen, X., Dui, W., Plemens, A., Khadr, L., et al. (2017). A germline-limited piggyBac transposase gene is required for precise excision in *Tetrahymena* genome rearrangement. Nucleic Acids Res. 45, 9481–9502.

89. Cervantes, M.D., Hamilton, E.P., Xiong, J., Lawson, M.J., Yuan, D., Hadjithomas, M., Miao, W., Orias, E., Cervantes, M.D., and Hamilton, E.P. (2013). Selecting one of several mating types through gene segment joining and deletion in *Tetrahymena thermophila*. PLoS Biol. 11, e1002284.

90. Li, H., Handsaker, B., Wysoker, A., Fennell, T., Ruan, J., Homer, N., Marth, G., Abecasis, G., Durbin, R., and Subgroup, G.P.D.P. (2009). The Sequence Alignment/Map format and SAMtools. Bioinformatics 25, 2078–2079.

91. Chaisson, M.J., and Tesler, G. (2012). Mapping single molecule sequencing reads using basic local alignment with successive refinement (BLASR): application and theory. BMC bioinformatics 13, 1–18.

92. Sheng, Y., Duan, L., Cheng, T., Qiao, Y., Stover, N.A., and Gao, S. (2020). The completed macronuclear genome of a model ciliate *Tetrahymena thermophila* and its application in genome scrambling and copy number analyses. Sci. China Life Sci. 63, 1534–1542.

93. Burger, G., Zhu, Y., Littlejohn, T.G., Greenwood, S.J., Schnare, M.N., Lang, B.F., and Gray, M.W. (2000). Complete sequence of the mitochondrial genome of *Tetrahymena pyriformis* and comparison with *Paramecium aurelia* mitochondrial DNA. J. Mol. Biol. 297, 365–380.

94. Hamilton, E.P., Kapusta, A., Huvos, P.E., Bidwell, S.L., Zafar, N., Tang, H., Hadjithomas, M., Krishnakumar, V., Badger, J.H., and Caler, E.V. (2016). Structure of the germline genome of *Tetrahymena thermophila* and relationship to the massively rearranged somatic genome. Elife 5, e19090.

95. Ye, F., Chen, X., Ju, A., Sheng, Y., Duan, L., Al-Rasheid, K.A.S., Stover, N.A., and Gao, S. (2024). Comprehensive genome annotation of the model ciliate *Tetrahymena thermophila* by in-depth epigenetic and transcriptomic profiling. bioRxiv, 2024.2001.2031.578305.

96. Ye, J., McGinnis, S., and Madden, T.L. (2006). BLAST: improvements for better sequence analysis. Nucleic Acids Res. 34, W6–W9.

97. Krueger, F., James, F., Ewels, P., Afyounian, E., and Schuster-Boeckler, B. (2021). FelixKrueger/TrimGalore: v0. 6.7-DOI via Zenodo. https://zenodo.org/record/5127899#.YofWf8NBzDc.

98. Kim, D., Paggi, J.M., Park, C., Bennett, C., and Salzberg, S.L. (2019). Graph-based genome alignment and genotyping with HISAT2 and HISAT-genotype. Nat. Biotechnol. 37, 907–915.

99. Liao, Y., Smyth, G.K., and Shi, W. (2014). featureCounts: an efficient general purpose program for assigning sequence reads to genomic features. Bioinformatics 30, 923–930.

100. Warnes, G.R., Bolker, B.M., Bonebakker, L., Gentleman, R., Liaw, W., Lumley, T., Maechler, M., Magnusson, A., Moeller, S., Schwartz, M.L., et al. (2015). gplots: Various R programming tools for plotting data. https://rdrr.io/cran/gplots/.

