## Supplemental figures for "Identification and characterization of the *de novo* methyltransferases for eukaryotic N^6^-methyladenine (6mA)"

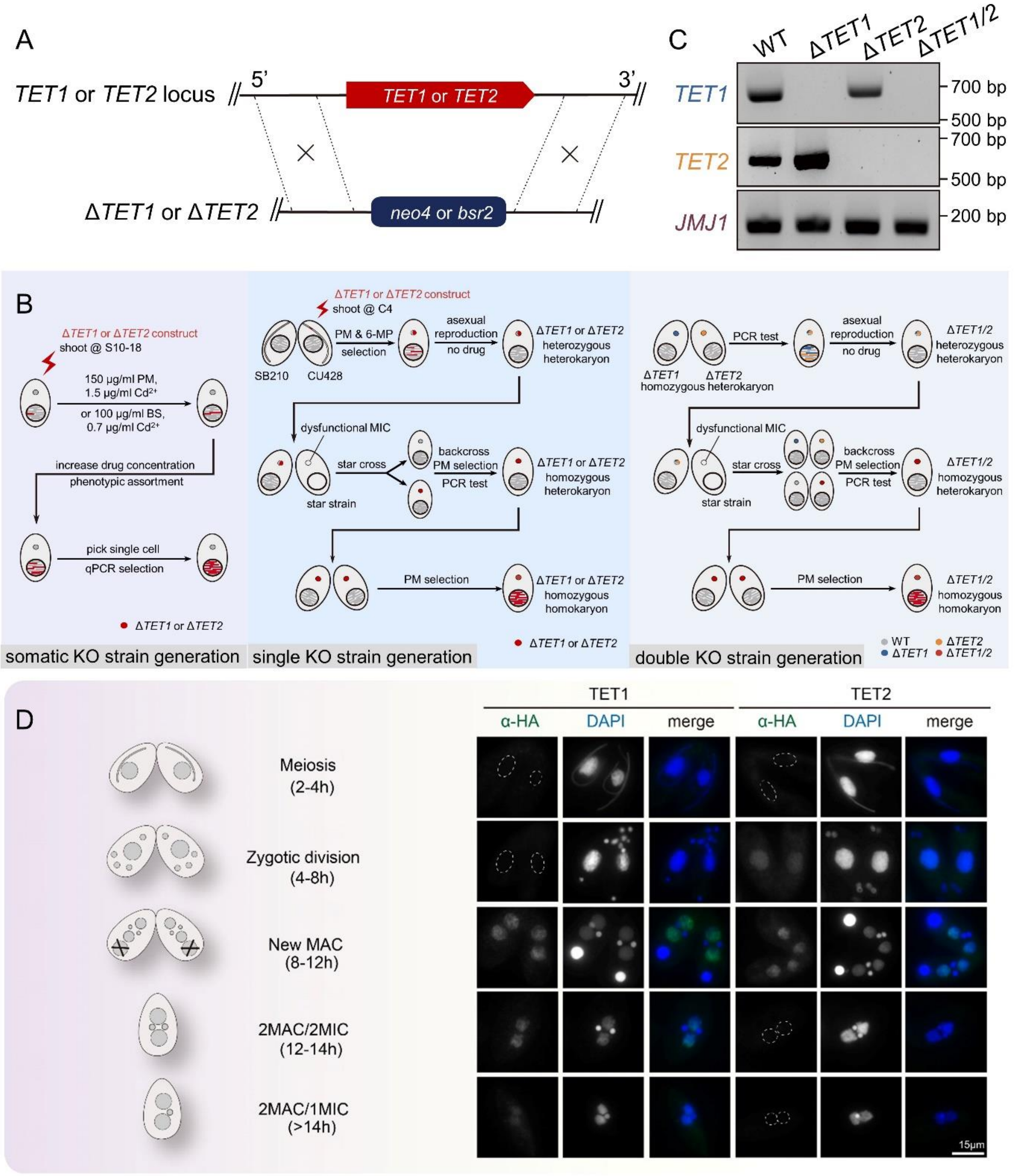


**Figure S1.** Generation and characterization of germline/somatic knockout (KO) strains.

A. Schematics showing the generation of Δ*TET1* and Δ*TET2* constructs. *TET1* or *TET2* coding sequences were replaced by the *neo4* or bsr2 cassette.

B. Schematics showing the generation of somatic and germline single and double KO cells. S10-18, 10-18h after starvation; C4, 4h post-mixing; PM, Paromomycin; BS, Blasticidin S; $Cd^{2+}$, $CdCl_2$.

C. RT-PCR analysis indicated that *TET1* and/or *TET2* were completely removed in germline KO cells. Total RNA was extracted from conjugative cells at 10h post-mixing and complementary DNA (cDNA) was used as the template for RT-PCR.

D. The distribution pattern of TET1 and TET2 along the conjugation progress, demonstrated by IF staining. Conjugative stages are distinguished by nuclear events (see schematics on the left). The outline of the nuclei without HA signal were delineated with dotted circles.

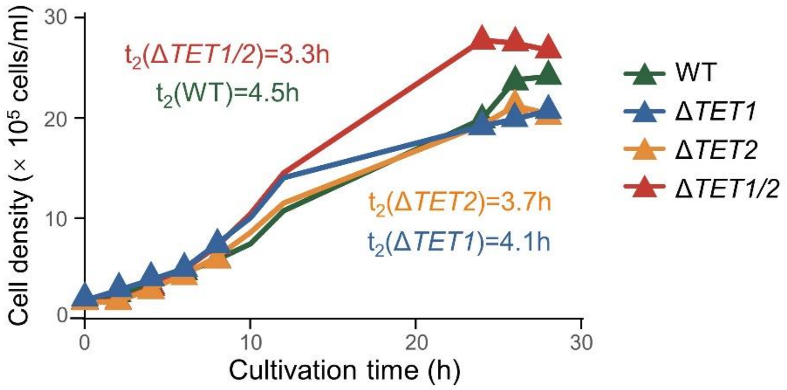


**Figure S2.** Loss of TET1 and/or TET2 did not affect the vegetative growth rate in KO cells. Cells were counted at the indicated time points. Doubling time (h) was calculated using the log-phase data. Three biological replicates were concurrently performed for each strain,  and the data were plotted using the mean values.

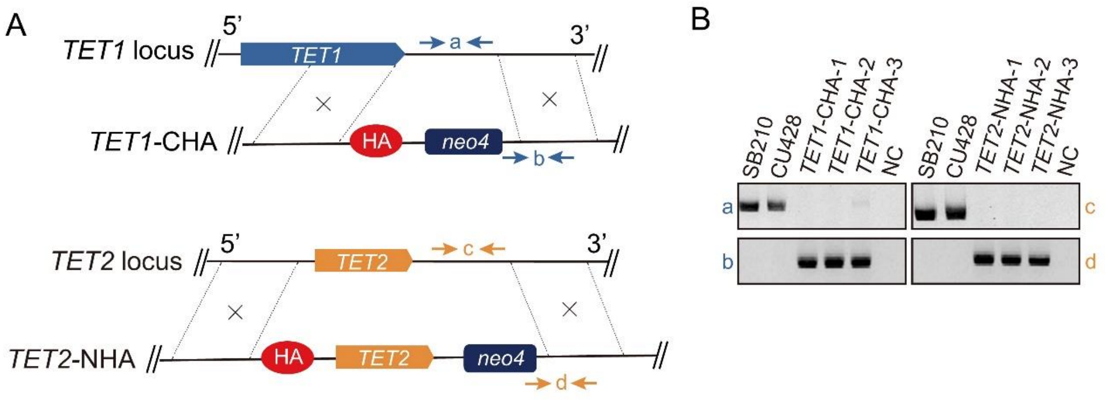


**Figure S3.** Generation and characterization of endogenously hemagglutinin (HA) tagged strains.

A. Schematics showing the generation of *TET1*-CHA-germline and *TET2*-NHA-soma constructs. Sequences coding for the hemagglutinin (HA) tag were inserted into the C-terminus of the *TET1* and the N-terminus of *TET2*, respectively. Arrows indicted the positions of testing primers in B.

B. PCR analysis confirmed that the endogenous loci of *TET1* and *TET2* were completely replaced by the assembled sequences. NC referred to the negative control for PCR.

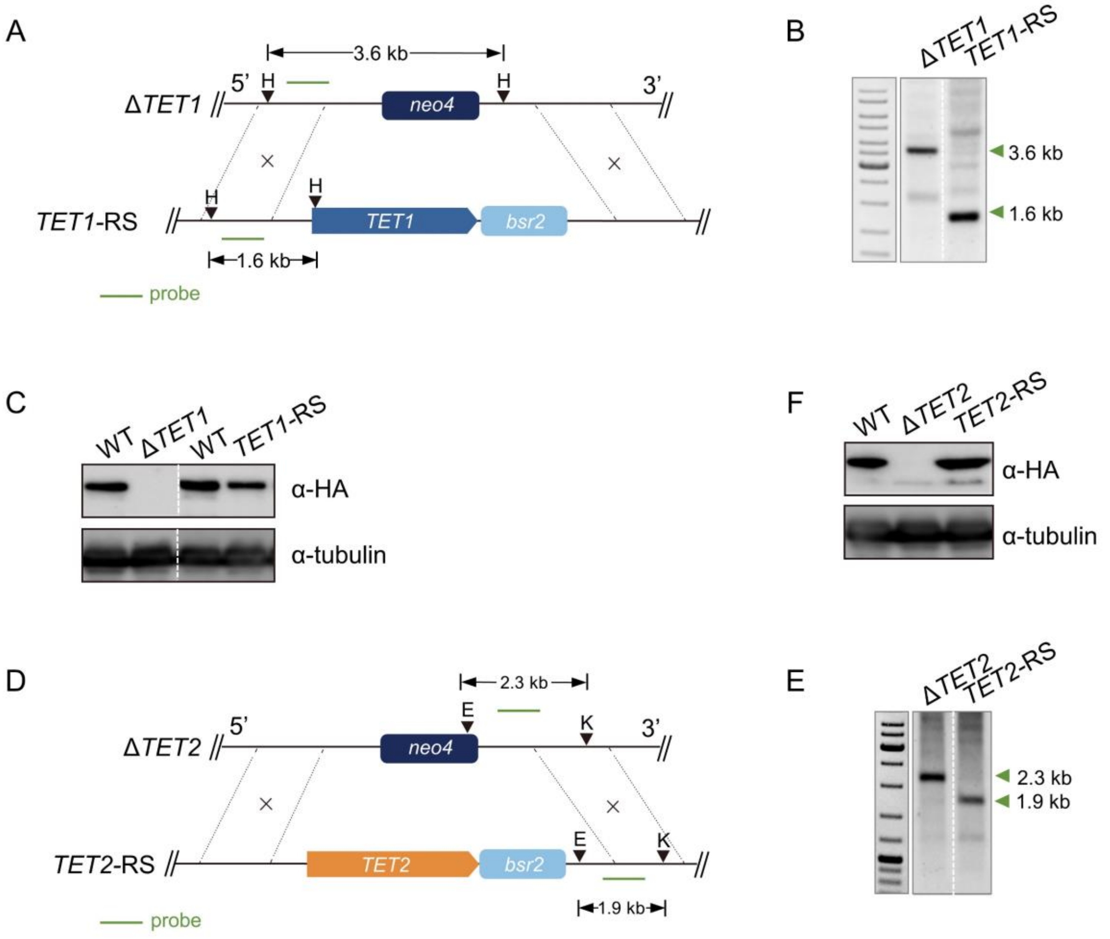


**Figure S4.** Generation and characterization of rescue strains.

A. Schematics showing the generation of *TET1* rescue constructs with WT (*TET1*-RS) copies. The *neo4* cassette in Δ*TET1* cells was replaced by the *TET1* full-length CDS, together with the *bsr2* cassette. H, *Hind*III restriction site. Green line, probe used for Southern blot assay.

B. The Southern blot assay confirmed that the *neo4* cassette in Δ*TET1* cells had been completely replaced by the assembled sequences of wild-type full-length CDS and the *bsr2* cassette. As shown in A, the probe used for detection is in a shared 5' region of *TET1* in WT and KO cells. *Hind*III was used to digest genomic DNA, producing different size of DNA fragments in KO and rescue cells. The Southern blot result showed that in rescue cells, the original sequence in the background KO cells had been fully replaced.

C. Western blot analysis revealed that the protein level of TET1 in *TET1* WT rescue cells were comparable to the level in WT cells. Cells were collected at 8h post-mixing, and proteins were precipitated using trichloroacetic acid (TCA). The expression of TET1 was detected using the anti-HA antibody (Roche, 3F10). Tubulin was detected as a control using the anti-α-tubulin antibody (Developmental

Studies Hybridoma Bank, 12G10).

D. Schematics showing the generation of *TET2* rescue construct (*TET2*-RS). The *neo4* cassette in Δ*TET2* cells was replaced by the assembled sequences of *bsr2* cassette and *TET2* full-length CDS. E, *EcoR*I restriction site. K, *Kpn*I restriction site. Green line, probe used for Southern blot assay.

E. The Southern blot assay confirmed that the *neo4* cassette in Δ*TET2* cells had been fully replaced by assembled sequences of *TET2* full-length CDS and the *bsr2* cassette. As shown in E, the probe used for detection is in a shared 3' region of *TET2* in WT and KO cells. *Sac*I *and BamH*I were used to digest genomic DNA, producing different size of DNA fragments in KO andrescue cells that contain the probe sequence. The Southern blot result showed that in rescue cells, the original sequence in the background KO cells had been fully replaced.

F. Western blot analysis revealed that the protein level of TET2 in *TET2* rescue cell was restored to the level in WT cells. The preparation of protein samples and the detection of expression levels were conducted according to the description in C.

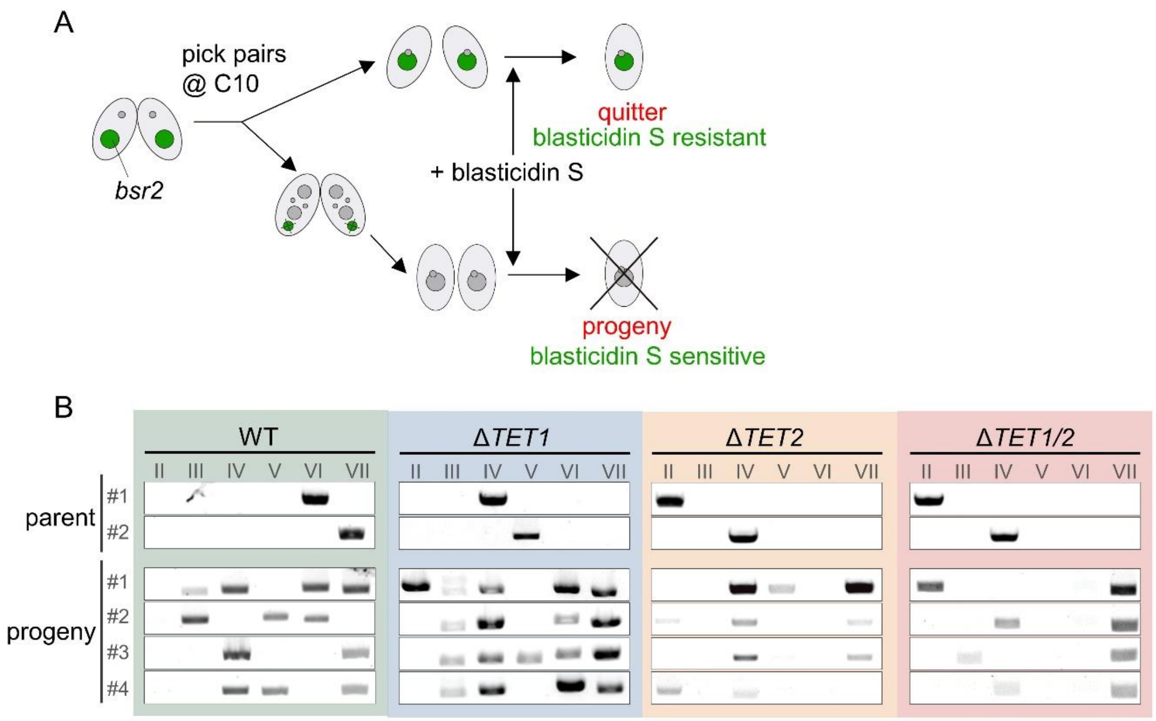


**Figure S5.** Selection of progenies.

A. Schematic diagram showing the process of selecting progenies. Cells of two different mating types were mixed and single pairs were hand-picked at 10h post-mixing. The blasticidin S resistance was tested to distinguish true progenies (sensitive) from quitters (resistant).

B. PCR amplification using mating type primers showed that progenies of the KO cells possessed different mating types from parental cells.

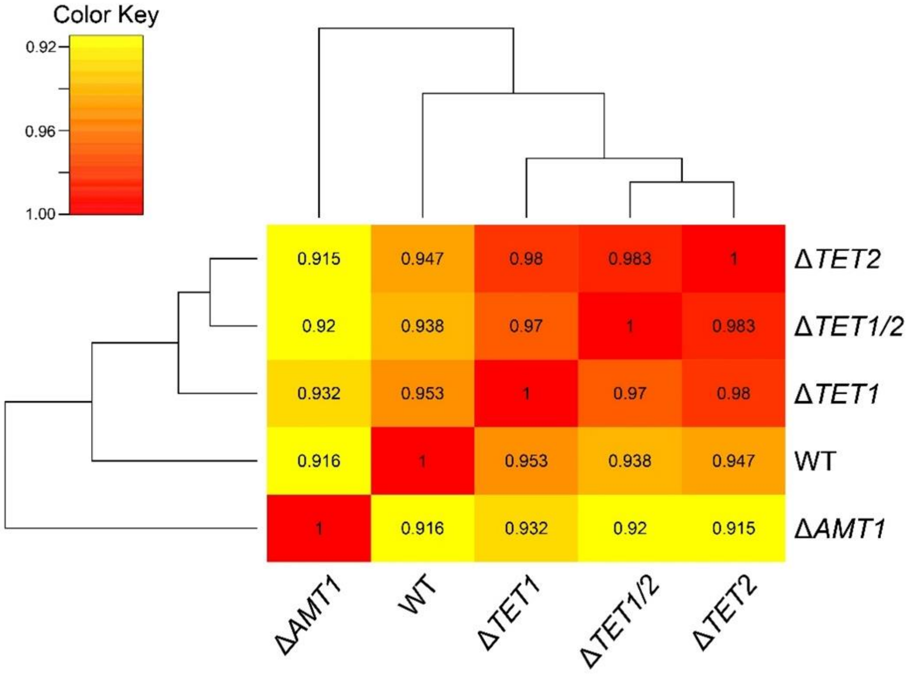


**Figure S6.** Comparison of gene expression profiles. Spearman's correlation coefficients were calculated for pair-wise comparison and visualized using a color scale. Note the similarities between KO cells (Δ*TET1*, Δ*TET2*, and Δ*TET1/2*) and their distinction from WT cells. Δ*AMT1* cells were used as an out-group.

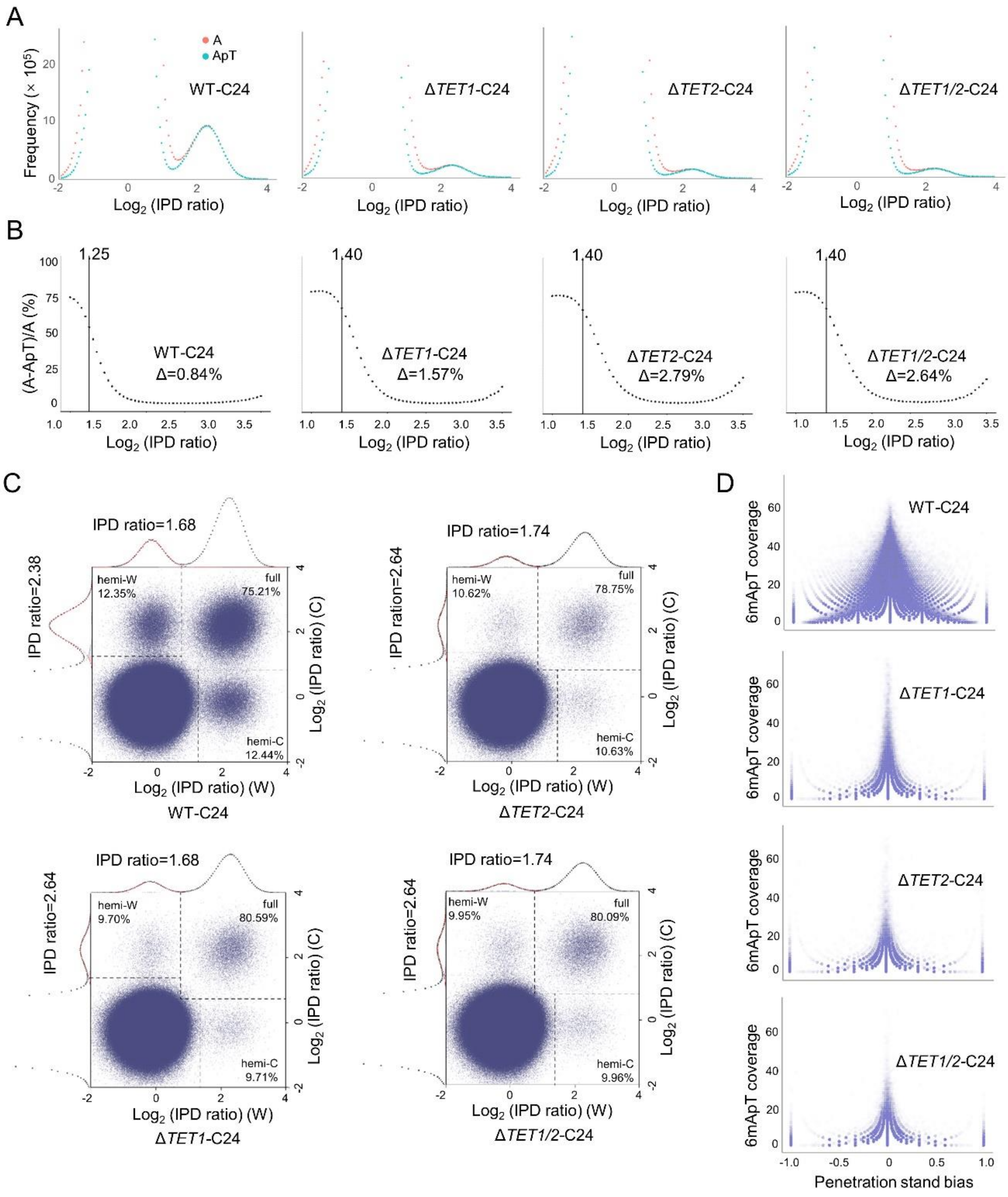


**Figure S7.** 6mA features in WT and KO cells at 24h post-mixing.

A. The distribution of IPD ratio was shown for all A sites (red) and A sites at the ApT dinucleotides (green). Despite a significant decrease of 6mA level in Δ*TET1,* Δ*TET2,* and Δ*TET1/2* cells, IPD ratios of adenines still maintained a bimodal distribution in KO cells, the same as that in WT cells.

B. Frequencies of four different dinucleotides (ApA, ApC, ApG, ApT) under different IPD ratios were counted and the frequency difference (Δ) between A and non-ApT sites was calculated, indicating the proportion of non-ApT sites relative to all A sites.

The minimum of the differential curve was marked reflecting the non-ApT/A ratio in 6mA, indicating the ApT preference in both KO and WT cells. Note that the ratio of non-ApT/A sites slightly increased after the deletion of *TET1* and/or *TET2*, which may be a combined effect of the overall decrease of 6mA level and the relatively constant sequencing background noise.

C. The plots showed the IPD ratios of each ApT dinucleotide on Watson and Crick strand. Different IPD ratio thresholds (shifted according to the bimodal distribution using Python script) were used to determine the methylation status of each ApT dinucleotide. The ratio of hemi- and full-methylated 6mApT positions appeared to be similar between KO and WT cells.

D. Penetration strand bias of each specific 6mApT position was defined as "$(W_{(penetration)}–C_{(penetration)})/(W_{(penetration)}+C_{(penetration)})$". The retained 6mApT positions in KO cells, especially those with higher 6mApT coverage, maintained a similar low penetration strand bias to WT cells.

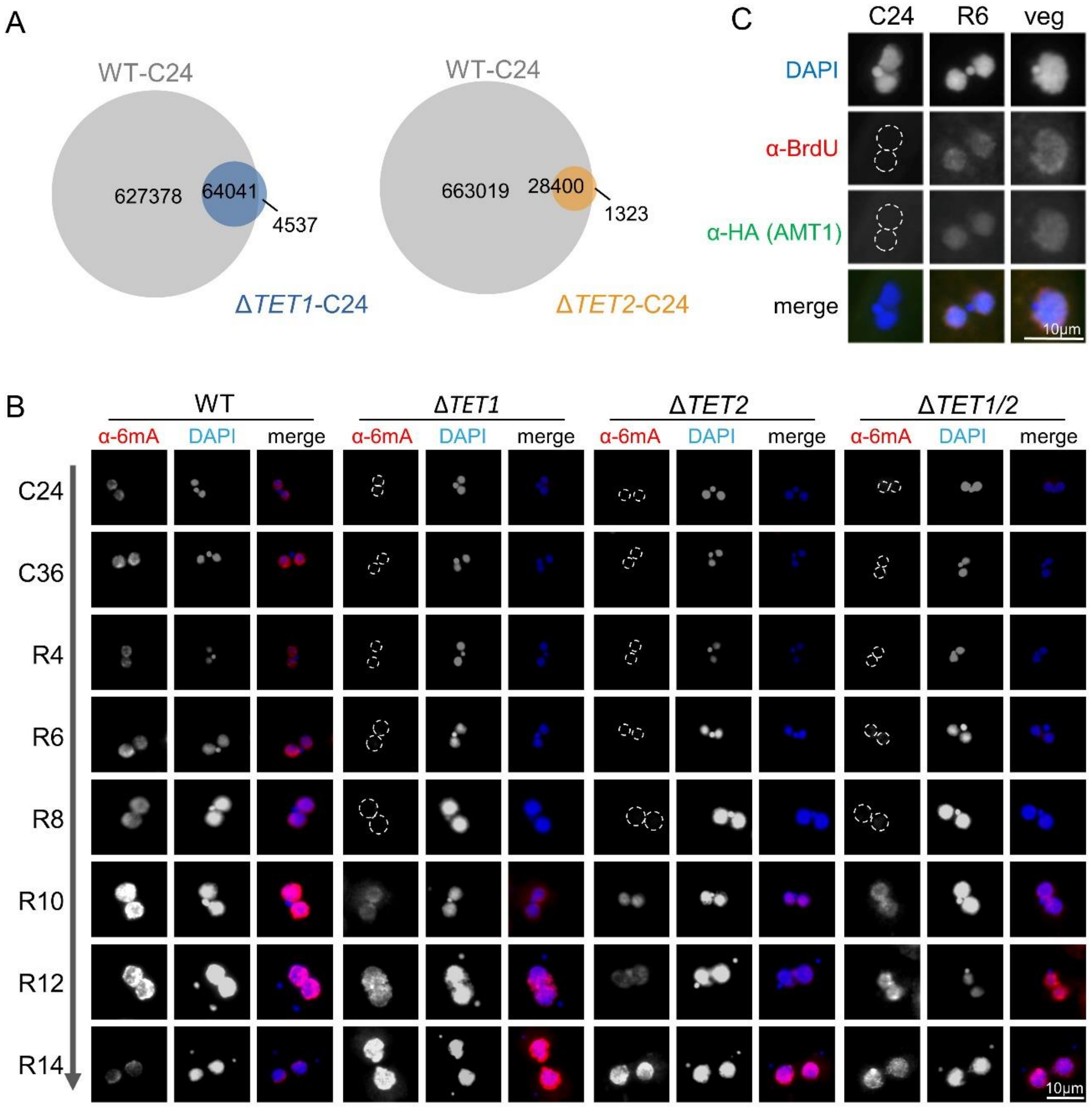


**Figure S8.** The upregulated protein level of AMT1 contributed to 6mA recovery in *TET1* and/or *TET2* KO cells.

A. Venn diagram showing the overlap between methylated 6mA sites in WT-C24 and Δ*TET1*-C24 and Δ*TET2*-C24. The vast majority of 6mApT positions in Δ*TET1* and Δ*TET2* cells were also methylated in WT cells. The same cutoff (6mApT coverage ≥ 10×) was applied to both WT and KO samples.

B. 6mA levels started to recover in KO cells after 10 hours of refeeding and reached a comparable level in KO and WT cells after 14 hours of refeeding, as shown by IF staining. The outline of the nuclei without 6mA signal were delineated with dotted circles.

C. The possible association between the AMT1 protein level and the occurrence of replication. Cells were labeled with 0.4mM BrdU for 30 min in dark and fixed at different time points: late conjugation 24h post-fixing (C24), 6h after refeeding (R6), and vegetative stage. The outline of the nuclei without BrdU or HA (AMT1) signal were delineated with dotted circles.
